# Isolation and characterization of rare circulating autoantibody-producing cells from patients with muscle-specific kinase myasthenia gravis

**DOI:** 10.1101/488247

**Authors:** Kazushiro Takata, Panos Stathopoulos, Michelangelo Cao, Marina Mané-Damas, Miriam Fichtner, Erik S. Benotti, Leslie Jacobson, Patrick Waters, Sarosh R. Irani, Pilar Martinez-Martinez, David Beeson, Mario Losen, Angela Vincent, Richard J. Nowak, Kevin C. O’Connor

## Abstract

Myasthenia gravis (MG) is a chronic autoimmune disorder characterized by muscle weakness and caused by autoantibodies that bind to functional membrane proteins at the neuromuscular junction. Most patients have autoantibodies to the acetylcholine receptor (AChR), but a subset of patients instead have autoantibodies to muscle specific tyrosine kinase (MuSK). MuSK is an essential component of the Agrin/Lipoprotein receptor-related protein 4(LRP4)/MuSK/downstream of tyrosine kinase 7 (DOK7) pathway that is responsible for synaptic differentiation, including clustering of AChRs at the neuromuscular junction, both during development and in adult muscle. Nerve-released Agrin binds to LRP4 which then binds to MuSK, stimulating autophosphorylation and recruitment of DOK7 to complete the membrane component of the pathway. Serum-derived IgG4 subclass MuSK autoantibodies prevent the binding of LRP4 to MuSK, subsequently impairing autophosphorylation, resulting in the loss of Agrin-induced AChR clustering in the mouse myotube-forming C2C12 line. Although this autoimmune mechanism appears well understood, MuSK autoantibodies from patients are polyclonal. Most are IgG4 but IgG1, 2 and 3 are also present and can achieve similar results on AChR clustering. In addition, most bind the first Ig-like domain in MuSK, however some patients harbor serum autoantibodies that recognize other domains in MuSK. We sought to establish individual MuSK IgG clones so that the disease mechanisms could be better understood. We isolated MuSK autoantibody-expressing B cells from MuSK MG patients with a fluorescent-tagged MuSK antigen multimer and generated a panel of human monoclonal autoantibodies (mAbs) from these cells. We produced 77 mAbs from single B cells collected from six MuSK MG patients. Here we focused on three highly specific mAbs that bound quantitatively to MuSK in solution, to MuSK-expressing HEK cells and at mouse neuromuscular junctions where they co-localized with AChRs. These three IgG isotype mAbs (two IgG_4_ and one IgG_3_ subclass) recognized the Ig-like domain 2 of MuSK. The three MuSK mAbs inhibited Agrin induced-AChR clustering in C2C12 myotubes, but intriguingly, they enhanced rather than inhibited MuSK phosphorylation. This approach for *ex vivo* isolation of MuSK MG autoantibody-producing cells and production of human recombinant mAbs has identified distinct autoantibody specificities and likely divergent effector mechanisms. Collectively, these findings will enable a better understanding of MuSK autoantibody-mediated pathology.

## Introduction

Patients with myasthenia gravis (MG) experience skeletal muscle weakness, worsened by activity. Typically, they present with ocular muscle weakness which then generalizes to involve limb muscles and, particularly, bulbar and respiratory muscles (1, 2). The molecular immunopathology of MG is directly attributed to the presence of circulating autoantibodies specifically targeting extracellular domains of functional postsynaptic membrane proteins at the neuromuscular junction (NMJ) (1, 3). The disease has multiple subtypes, defined by different autoantibody targets (4-7). Autoantibodies to the acetylcholine receptor (AChR), present in around 85% of patients, are mainly IgG1 and cause loss of AChRs via divalent binding, leading to internalization of AChRs and complement-mediated damage to the NMJ (1, 8). Some of the 15% of patients without AChR autoantibodies have, instead, autoantibodies to muscle-specific kinase (MuSK) (7) or less commonly Lipoprotein receptor-related protein 4 (LRP4) (9, 10). The MuSK autoantibody form of MG can be severe as it usually involves mainly bulbar muscles (11), affecting speaking, chewing, swallowing, and breathing, and can cause permanent muscle atrophy over time (12, 13). MuSK autoantibodies are particularly interesting because they are predominantly (14) of the non-complement activating IgG4 subclass; a subclass that can be functionally monovalent for antigen binding and hence does not cross-link its antigen (15). Yet MuSK autoantibodies are demonstrably pathogenic, which can be established by passive transfer of the human disease phenotype to mice by injection of patients’ IgG, or active immunization with MuSK (16-18).

MuSK is an essential component of the Agrin/ LRP4/MuSK/downstream of tyrosine kinase 7 (DOK7) pathway that is responsible for clustering of AChRs at the NMJ, both during development and in mature muscle (19). Serum-derived MuSK autoantibodies recognize mainly the N-terminal Ig-like domain 1 of MuSK, and prevent the binding of LRP4 to MuSK (20-22). As a result, autophosphorylation of MuSK is inhibited and DOK7 is not recruited to complete the pathway. These effects can be demonstrated in the mouse myotube-forming C2C12 cell line, where MuSK autoantibodies prevent Agrin/LRP4-induced clustering of AChRs. In this model, isolated antigen binding fragments (Fabs) from MuSK-specific antibodies are sufficient to inhibit AChR clustering (23). By contrast, AChR autoantibodies require divalent binding in order to cause loss of AChRs (8, 24, 25). Although some of the mechanisms underlying MuSK-autoantibody-associated MG appear well understood, patients’ autoantibodies are heterogeneous. For instance, IgG1, 2 and 3 MuSK autoantibodies exist in most patients and their pathogenic mechanisms have not been well studied. Moreover, it is unclear whether autoantibodies to domains other than the first Ig-like domain in MuSK may contribute to disease. We sought to establish individual MuSK IgG clones so that the mechanisms in this disease could be better analyzed both *in vitro* and *in vivo*.

All forms of MG improve with immunotherapies, but B cell depletion with a therapeutic monoclonal antibody (rituximab) against the B cell marker CD20 leads to substantial reductions in MuSK autoantibodies and relatively quick clinical improvement (11, 26, 27). The success of anti-CD20 therapy suggests that the autoantibodies are derived from circulating MuSK-specific B cells rather than bone marrow resident long-lived plasma cells (LLPCs). LLPCs, which produce the vast majority of circulating antibody, express negligible levels of CD20 and thus are not targets of rituximab treatment (28). This is confirmed by unchanged serum immunoglobulin and vaccine-specific titers post-treatment (26, 29, 30). Accordingly, we proposed a speculative model in which an autoreactive fraction of memory B cells and circulating short-lived plasmablasts are responsible for much of the MuSK autoantibody production (3, 31) and recently demonstrated that circulating plasmablasts do indeed contribute to MuSK MG autoantibody production (32).

Given the accessibility of circulating autoantibody-producing cells, we adapted a previously reported approach (33) to produce a high-avidity fluorescent-tagged MuSK tetramer that could identify and assist in sorting rare autoantibody-expressing B cells from patient-derived blood samples. The specificity of the isolation was validated by single-cell sorting of antigen-labeled B cells and by reconstruction of recombinant human monoclonal antibodies, which were then tested for binding to MuSK.

## Methods

### Study Approval

This study was approved by the Human Investigation Committee at the Yale School of Medicine. Informed consent was obtained from all subjects.

### Isolation of serum and PBMC from MuSK MG patients

Peripheral blood samples were obtained from two healthy donors (HD1-2) and six patients (MuSK1-6) with autoantibody and clinically confirmed MuSK MG. Patients showed typical clinical and serological features of MuSK MG (**Table 1**). PBMCs were isolated by Ficoll separation and stored in liquid nitrogen until use, using a described protocol (34). Time-locked serum specimens were also obtained.

**Table 1.** Study subject clinical, laboratory and demographic data.

| Patient | Sex | Age at TOC | Diagnosis | Ab status | MGFA class | Rituximab | Other therapy | Serum MuSK Ab titer |
| --- | --- | --- | --- | --- | --- | --- | --- | --- |
| MuSK1 | F | 37 | Generalized MuSK MG | MuSK | IIIB | 1cycle<sup>A</sup> 19 mo before TOC | None | 5.7<sup>B</sup> |
| MuSK2a | F | 45 | Generalized MuSK MG | MuSK | I | 4 cycles 78 mo before TOC | None | 11.6<sup>B</sup> |
| MuSK3 | F | 53 | Generalized MuSK MG | MuSK | IIA | 2cycles 21 mo before TOC | prednisone 10mg/d PLEX | 0.9<sup>B</sup> |
| MuSK4 | F | 35 | Generalized MuSK MG | MuSK | IIB | None | mycophenolate 1500mg/day | 1:1280<sup>C</sup> |
| MuSK5 | F | 31 | Generalized MuSK MG | MuSK | IIB | None | PLEX | 1:640<sup>C</sup> |
| MuSK6 | F | 63 | Generalized MuSK MG | MuSK | IIB | 5 cycles 20 mo before TOC | None | 1:5120<sup>C</sup> |
| HD1 | M | 37 | - | - | - | - | - | - |
| HD2 | F | 30 | - | - | - | - | - | - |

<sup>A</sup>A cycle of rituximab consisted of 1 infusion per week for 4 weeks; dose per infusion: 375 mg/m<sup>2</sup>.
<sup>B</sup>Determined by radioimmunoassay; levels in nmol/l. <sup>C</sup>Determined by RIA; levels reported as serum titer.
F, female; M male; HD, healthy donor; MGFA, MG Foundation of America clinical classification; MuSK, muscle-specific tyrosine kinase; NA, not available; PLEX, plasma exchange; TOC, time of collection; mo, month. Additional specimens from three of the patients (MuSK1, MuSK2a and MuSK3) had been studied in our previous investigation and are named as indicated in that report (32).

### MuSK multimer generation

The extracellular domain of MuSK was cloned into the pMT/Bip/His-A vector. The C-terminal region contained a short, flexible linker followed by the a BirA site (amino acids: GLNDIFEAQKIEWHE), downstream of which was a thrombin-cleavable (amino acids: LVPRGS) 6x histidine-tag. Protein expression was induced in S2 *Drosophila* cells. Culture supernatant was collected and MuSK protein was purified using cobalt resin beads (Thermo scientific) according to the manufacturer’s instructions. For tetramer formation, MuSK protein was biotinylated by incubation with BirA enzyme at a 1:100 molar ratio overnight at 4 °C in a buffer containing 50 mM tris, 50 mM bicine pH 8.3, 10 mM magnesium acetate, 10 mM adenosine-5’-triphosphate and 50 μM biotin. Excess biotin was removed using a 10kDa MWCO Slide-A-Lyzer dialysis cassette (Thermo scientific). Fluorescent multimers were formed using stepwise addition of allophycocyanin (APC)-conjugated streptavidin (Invitrogen) to biotinylated MuSK until a 1:4 molar ratio was reached.

### Flow cytometry and cell sorting

For sorting MuSK multimer reactive B cells, B cells were enriched from PBMCs using negative-selection beads (Stemcell Technologies). They were incubated with live/dead stain, then stained with 20 μg/ml MuSK multimer on ice for 30 min. Cells were then co-stained (using manufacturer’s recommended dilutions) with fluorescently labeled antibodies against CD3 (Invitrogen, Pacific orange; UCTH1), CD14 (Invitrogen, Pacific orange; TUK4), CD19 (BD Biosciences, PE Cy7; SJ25C1), CD27 (BD Biosciences, PE; M-T271), and CD38 (BD Biosciences, V450; HB7) before sorting on a FACSaria (BD Biosciences) instrument. For the purposes of general B cell immunophenotyping, B cells were defined as live, CD3^-^, CD14^-^, CD19^+^ cells; memory B cells as live, CD3^-^, CD14^-^, CD19^+^, CD27^+^, CD38^-^ cells and antibody secreting cells (plasmablasts) as CD3^-^, CD14^-^, CD19^+^, CD27^+^, CD38^high^.

### MuSK, AChR and myelin oligodendrocyte glycoprotein (MOG) monoclonal antibodies

A set of monoclonal antibodies for binding to either MuSK, AChR and MOG were used as controls. Cell culture supernatant from either an established murine MuSK mAb (4A3)(32) or a murine MOG mAb (8-18C5) (35) hybridoma was applied to a protein G-Sepharose column in order to isolate the IgG. We also engineered the MuSK mAb (4A3) and the MOG mAb (8-18C5) as chimeric mouse-human recombinant mAbs. They were produced to contain the murine mAb H- and L-chain V regions fused to the respective human constant regions using an approach we described (36). These chimeric recombinant mAbs served as positive controls in the human antibody binding assays and did not require a substitute (murine specific) secondary antibody because the constant regions were identical to those of human mAbs. The AChR mAb (637) was derived from a human MG thymus (24, 25, 37). The variable regions were synthesized as gBlock gene fragments (Integrated DNA Technologies) then subcloned into expression vectors, expressed and purified using approaches we described ((38) and Cotzomi, Stathopoulos *et al.* submitted).

### Recombinant human monoclonal antibody production, IgG subclass determination and sub-cloning

Reverse transcription of fresh or frozen single cell-sorted, antigen-labeled B cell RNA, nested PCR reactions, cloning into IgG expression vectors, antibody expression, and purification were all performed as described (34). To determine the isotype and IgG subclass, a specialized single-cell PCR using left-over cDNA from the same single cells used to make individual mAbs, was performed. This PCR used a primer in a down-stream region of the IgG such that the PCR product included a region of the IgG 1, 2, 3, and 4 that is unique to each subclass (39, 40). Thus, identification of IgG subclass required sequence alignment of this region to each of the four human IgG subclasses. Individual pcDNA3.3-based expression vectors that included each of the human IgG subclass constant regions were constructed in-house using an approach we described (38). Following the determination of the endogenous IgG subclass for each human mAb, the variable heavy chain region was sub-cloned into the respective subclass-containing expression vectors.

### Cell-based antibody assays

Antibody binding was tested using live 293T HEK cells transiently transfected with DNA encoding either MuSK, AChR with Rapsyn or MOG using an assay protocol we described (32). The following plasmid constructs were used for expression: Full-length human MuSK (21) cloned into a pIRES2-EGFP vector, which delivered translation of MuSK and GFP separately. Human AChR α-, β-, δ- or ε-subunits cloned into pcDNA3.1-hygro plasmid vectors (41) and Rapsyn (41), which was cloned into a pEGFP-N plasmid. Full-length human MOG (42) was also cloned, like Rapsyn, into the pEGFP-N plasmid, which produced rapsyn or MOG as fusion proteins with C-terminal GFP. Cell-based assay (CBA) results were calculated as ΔMFI (mean fluorescence intensity) and percent of transfected cells that bound secondary antibody (termed % positive) as follows: (i) ΔMFI = Alexa Fluor 647 MFI in MuSK-GFP-transfected cells minus Alexa Fluor 647 MFI in untransfected cells of the same tube, (ii) % positive cells=% of cells in upper right quadrant / % of cells in upper right and lower right quadrant.

### MuSK protein mutagenesis

The generation of MuSK deletional mutants was performed by modifying the full-length MuSK expression construct. Regions coding for specific domains were deleted using the Q5 Site-Directed Mutagenesis Kit (BioLabs) according to the manufacturer’s instructions. Primer sequences were generated using the NEBaseChanger tool. All construct modifications were confirmed through Sanger-based DNA sequencing.

### Immunoglobulin sequence analysis

The heavy- and light-chain variable region germline gene segments were identified with the IMGT/V-QUEST program (43) version: 3.4.14 (10 September 2018) - IMGT/V-QUEST reference directory release: 201839-3 (26 September 2018). Somatic mutations resulting in replacement amino acids were evaluated through the alignment to germline genes provided by the IMGT/V-QUEST program. Immunoglobulin isotype and IgG subclasses were determined by aligning acquired sequences to those present in the IMGT repertoire reference set (44).

### Immunofluorescence mouse muscle sections

The binding of the different human mAbs was analyzed by immunofluorescence using cryosections of mouse tibialis anterior muscles. Muscles were cut longitudinally at 10?µm thickness on a Leica CM3050 S cryostat; sections were mounted on gelatin-coated glass slides and stored at −80 °C. After thawing, cryosections were fixed with 4% PFA for 10 min at room temperature and then blocked for 30 min with 2% bovine serum albumin. Sections were incubated for 1 hr at room temperature with one of the different human mAbs (1.5 µg/mL each), combined with Alexa-647-conjugated alpha-bungarotoxin (1:300, B35450, Thermo Fischer Scientific) and biotinylated mouse anti-SV2 (against synaptic vesicle glycoprotein 2A, 1 µg/mL, DS Hybridoma Bank). As controls, protein G purified IgG from a MuSK MG patient (final concentration 5 µg/mL) was used instead of the mAbs. After washing, slides were incubated with human Fcγ-specific Alexa-488-conjugated goat F(ab’)_2_ (3 µg/mL, 109-546-170, Jackson ImmunoResearch), combined with Alexa-594-conjugated streptavidin (1:20,000, S11227, Invitrogen) and Hoechst 33342 (2 µg/mL; B2261, Sigma) for 1 hr at room temperature in the dark. Sections were washed and mounted with 80% glycerol. All washing steps consisted of three incubation of the slides (5 min at room temperature) in 0.05% Triton-X100. Endplates were identified using the red immunofluorescence of the alpha-bungarotoxin staining. Triple fluorescent photomicrographs of the endplate regions were acquired using µManager software ver2.0 (45) on an Olympus BX51WI spinning disk confocal fluorescence microscope with a Hamamatsu EM-CCD C9100 digital camera. Endplates were analyzed using ImageJ software as described (45, 46). All staining procedures and fluorescent analysis were performed on coded samples by two independent, blinded investigators.

### AChR clustering assay

The AChR clustering assay was performed as described (7). Briefly, C2C12 mouse myoblasts (ATCC) were grown in DMEM supplemented with 20% fetal bovine serum (FBS) and 1% penicillin/streptomycin (Gibco). C2C12 cells were plated in 24-well plates and differentiated with DMEM supplemented with 2% FBS, 0.5% penicillin/streptomycin and 1 μM insulin (Sigma). As soon as fusion was evident (36 to 48 hr), AChR clustering was induced for 14 hr with 10 ng/mL Agrin (R&D Systems). Monoclonal antibodies were applied at 1 μg/mL together with Agrin or alone. After the induction of AChR clustering, AChRs were visualized through the application of 1 μg/mL Alexa 647-labelled alpha-bungarotoxin (Invitrogen) for 1 hr at 37 °C. Following staining, cells were washed twice with medium (5 min at 37° C) and fixed with 3% PFA for 20 min at room temperature. Microscopy was performed at a 100X magnification on a Leica DMi8 fluorescence microscope. For each well, four visual fields of 100% myotube confluence were selected on phase contrast and captured on fluorescence; AChR clusters were counted using ImageJ software. For each condition duplicate wells were used and the mean of the clusters per visual field per condition were calculated. Experiments were performed at a minimum of three repetitions and were normalized for the effect of Agrin. Reported results are from experiments in which a minimum three-fold effect of Agrin-induced clustering over the baseline was observed.

### MuSK tyrosine phosphorylation assays

Myotubes (C2C12) were stimulated with 0.4 nM neural Agrin or Agrin together with IgG from MuSK MG patients, or mAbs for 30 min at 37 °C. To detect and quantify MuSK phosphorylation levels, myotubes were extracted in cold lysis buffer (10 mM Tris-HCl, 1 mM EDTA, 100 mM NaCl, 1% Triton X-100, 1x protease inhibitor cocktail, 1x phosphatase inhibitor cocktail) followed by centrifugation. To precipitate endogenous MuSK, the whole cell lysate was incubated with an anti-MuSK polyclonal antibody (AF562, R&D Systems) at 4 °C overnight. Bound antibody was captured with Dynabeads protein G (Invitrogen). Bead-precipitated proteins were eluted into SDS sample buffer, subjected to SDS-PAGE and incubated with monoclonal mouse anti-phosphotyrosine antibody (4G10, Upstate Biotechnology). The membrane was then harshly stripped (in 62.5 mM Tris buffer pH 6.8, containing 2.0% SDS and 0.8% β-mercaptoethanol) and re-probed for MuSK by incubating with a goat anti-MuSK polyclonal antibody (AF562, R&D). Densitometry of bands was analyzed using ImageJ software and the level of MuSK phosphorylation normalized to levels of immunoprecipitated MuSK.

### Statistics

AChR clustering on the C2C12 cells and MuSK tyrosine phosphorylation were analyzed using a one-way ANOVA with Dunnett’s correction. Statistics were performed on GraphPad Prism (version 7.0a) software.

## Results

### Study subjects

Patients (n=6, all female, mean age 44 ± 12 years (range 37–63), consistent with reported (47-49) demographics of MuSK MG) with laboratory and clinically confirmed MuSK autoantibody-positive MG were selected for study. Their clinical severity and serum autoantibody status at the time of sampling are summarized in **Table 1**. The controls included two healthy individuals, one male aged 37 years and one female aged 30 years (**Table 1**). Both had no history of autoimmune disease and no recent inflammatory events and were negative for serum MuSK autoantibodies.

### Generation of a multimeric, fluorescent MuSK antigen

We expressed the extracellular domain of MuSK, which was tagged, at the C-terminus, with a BirA site that allows for post-translational biotinylation. The addition of allophycocyanin (APC)-conjugated streptavidin then was used to generate a fluorescent antigen tetramer/multimer **(Figure S1A).** To validate that MuSK-specific antibodies were able to recognize the tetramer, antibody binding was tested in a flow cytometry-based assay. Flow cytometry beads, coated with anti-mouse immunoglobulin antibodies, were incubated with either the hybridoma-derived murine mAb, 4A3, that recognizes human MuSK or a control mAb, 8-18C5, that recognizes human myelin oligodendrocyte glycoprotein (MOG). Antibody-coated beads were incubated with fluorescent MuSK multimers and then analyzed by flow cytometry. The MuSK multimer was bound by beads that were coated with the MuSK specific mAb, but not those coated with the MOG mAb **(Figure S1B)**. These data established that the multimerized, labeled MuSK retained properties required for antibody recognition and was suitable for identifying B cells expressing MuSK-specific B cell receptors (BCR).

### Identification and isolation of MuSK-specific B cells using labelled antigen

We recently demonstrated (32) that MuSK autoantibody-producing cells can be identified in the circulation. Thus, we utilized the multimerized, labeled MuSK to enrich for such antigen-specific B cells from PBMCs. Using flow cytometry, purified B cell populations were separated into two populations; antibody secreting cells (plasmablast-like phenotype CD19^+^CD27^+^CD38^high^) and antigen-experienced B cells (memory-like phenotype CD19^+^CD27^+^CD38^-^). Cells within those two populations that bound the fluorescent MuSK multimer (**Figure S2**) were deposited as single cells into the wells of PCR plates.

### Screening of recombinant human monoclonal antibodies

We cloned and expressed human recombinant mAbs from single sorted B cells or plasmablasts, which were positive for staining with the fluorescent MuSK multimer. We cloned 77 mAbs from the six MuSK MG patients, which were derived from memory B cells or plasmablasts that recognized MuSK, and another 29 from the two healthy controls. For initial antigen specificity screening, all of the variable heavy chain domains were cloned into a human IgG1 subclass expression vector, irrespective of their native isotype or IgG subclass usage, which was not determined at this step. The variable light chain domains were cloned into either a kappa or lambda expression vector based on their native usage. We first screened the mAbs for MuSK binding at 10 μg/mL using a live cell-based assay (CBA). At this mAb concentration, many of the mAbs, including those from the healthy controls, showed binding. However, using a concentration of 1μg/mL, three mAbs (MuSK1A, MuSK1B and MuSK3B) from two patients (MuSK1 and MuSK3), unlike the mAbs from the two healthy donors (HD), demonstrated robust binding to MuSK (**Figure 1, Table 2**). Most other mAbs from the MuSK patients (MuSK2a, MuSK4, MuSK5 and MuSK6) did not bind at this concentration (**Figure 1**). Consequently, we focused on the three robust-binding mAbs (MuSK1A, MuSK1B and MuSK3B) along with an additional robust-binding MuSK mAb that we had previously isolated (MuSK3-28) (32).

### Cellular origins and binding properties of MuSK-specific recombinant monoclonal antibodies

During the MuSK-specific B cell sort, the FACS analyzer marked each cell in the scatter plot and its corresponding position in the 96 well plate (index-sorting). After the mAbs were expressed and their MuSK specificity validated on the CBA, the exact position of the cell on the scatter plot was determined and consequently its phenotype was assessed. With this approach, we determined (**Table 2**) that mAbs MuSK1A and MuSK3B were derived from B cells displaying a memory-like phenotype (CD19^+^CD27^+^CD38^-^) and MuSK1B was derived from a B cell displaying a CD38+ plasmablast-like phenotype (CD19^+^CD27^+^CD38^high^). The mAb MuSK3-28 was isolated from a single-cell sorted total plasmablast population (32), without the use of the MuSK multimeric antigen.

**Table 2.**
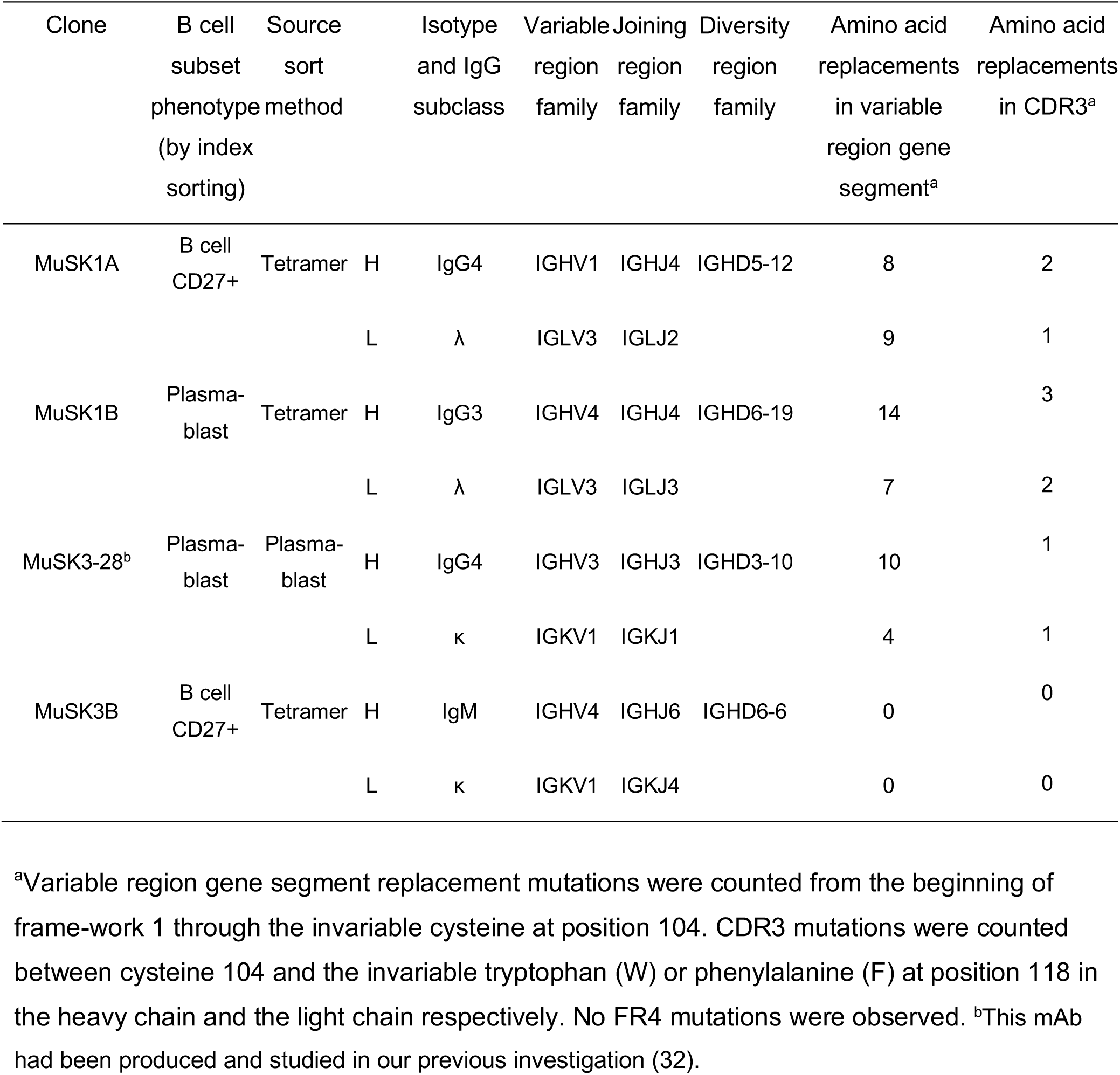
Molecular characteristics of MuSK-binding human recombinant monoclonal autoantibodies.

| Clone | B cell subset phenotype (by index sorting) | Source sort method |  | Isotype and IgG subclass | Variable region family | Joining region family | Diversity region family | Amino acid replacements in variable region gene segment<sup>a</sup> | Amino acid replacements in CDR3<sup>a</sup> |
| --- | --- | --- | --- | --- | --- | --- | --- | --- | --- |
| MuSK1A | B cell CD27+ | Tetramer | H | IgG4 | IGHV1 | IGHJ4 | IGHD5-12 | 8 | 2 |
|  |  |  | L | λ | IGLV3 | IGLJ2 |  | 9 | 1 |
| MuSK1B | Plasma-blast | Tetramer | H | IgG3 | IGHV4 | IGHJ4 | IGHD6-19 | 14 | 3 |
|  |  |  | L | λ | IGLV3 | IGLJ3 |  | 7 | 2 |
| MuSK3-28<sup>b</sup> | Plasma-blast | Plasma-blast | H | IgG4 | IGHV3 | IGHJ3 | IGHD3-10 | 10 | 1 |
|  |  |  | L | κ | IGKV1 | IGKJ1 |  | 4 | 1 |
| MuSK3B | B cell CD27+ | Tetramer | H | IgM | IGHV4 | IGHJ6 | IGHD6-6 | 0 | 0 |
|  |  |  | L | κ | IGKV1 | IGKJ4 |  | 0 | 0 |

<sup>a</sup>Variable region gene segment replacement mutations were counted from the beginning of frame-work 1 through the invariable cysteine at position 104. CDR3 mutations were counted between cysteine 104 and the invariable tryptophan (W) or phenylalanine (F) at position 118 in the heavy chain and the light chain respectively. No FR4 mutations were observed. <sup>b</sup>This mAb had been produced and studied in our previous investigation (32).

**Figure 1.**
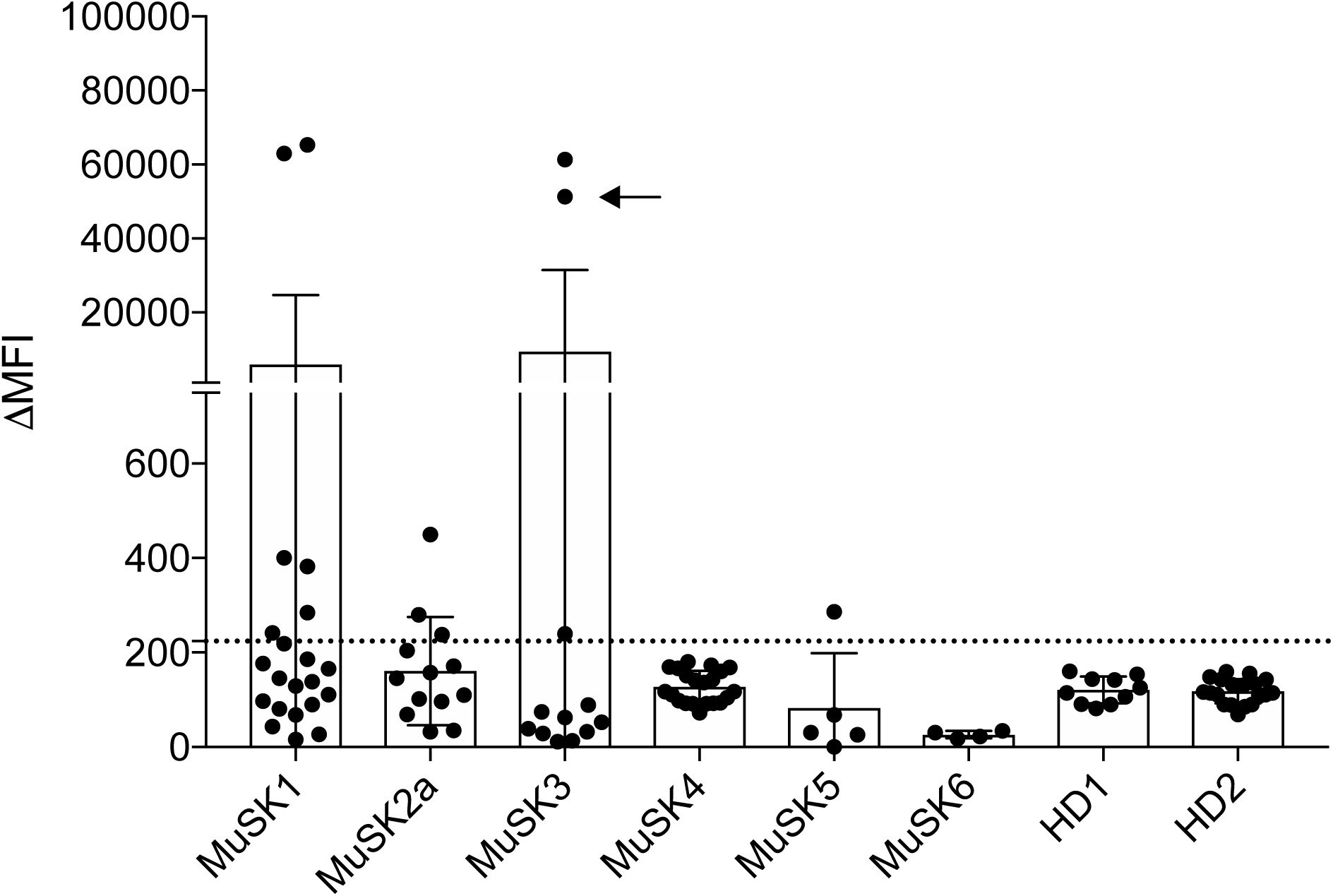
Screening of human recombinant monoclonal antibodies. Recombinant mAbs were produced from single MuSK multimer-sorted B cells. Binding of these clones to MuSK-expressing cells was determined using a flow cytometry-based antibody binding assay. Each data point represents the mean ΔMFI of each mAb tested at 1μg/mL in triplicate. Bars represent the mean of means and error bars SDs. The mAbs were derived from patients with MuSK MG and healthy donors (HD): MuSK1 (n=22), MuSK2a (n=5), MuSK3 (n=12), MuSK4 (n=13), MuSK5 (n=21), MuSK6 (n=6), HD1 (n=10) and HD2 (n=19). A human recombinant MuSK mAb that we previously produced from single-cell sorted plasmablasts (indicated with an arrow) was included with those tested from patient MuSK3. Values greater than the mean + 4SD of the HD-derived mAbs (indicated by the horizontal dotted line) were considered positive.

### Isotype, IgG subclass usage and molecular properties of MuSK-specific recombinant monoclonal antibodies

The native isotype and IgG subclass was determined using an additional PCR and sequencing step. The IgG4 subclass was used by the B cell that expressed mAb MuSK1A (**Table 2**), mAb MuSK1B was natively expressed using IgG3 and mAb MuSK3B was IgM (**Table 2)**. The mAb MuSK3-28 was natively expressed using IgG4. We next examined the B cell receptor characteristics of the MuSK-specific autoantibodies. Sequence analysis of the four MuSK mAbs revealed that these autoantibodies are represented by diverse clones that utilize different variable region gene segments (**Table 2**). A number of somatic mutations, a hallmark of affinity maturation, had accumulated in the variable heavy (VH) and variable light (VL) CDR regions of the three IgG isotype mAbs, strongly suggesting that antigenic selection had occurred. The fourth mAb which used IgM, did not include any somatic hypermutations. These immunoglobulin sequencing data show that MuSK autoantibodies are mostly class switched and suggest that the development of MuSK autoantibodies often requires the processes of clonal selection affinity maturation and class switching.

### Binding properties of MuSK-specific recombinant monoclonal antibodies

We next sought to examine the binding properties of the IgG mAbs. The IgM isotype-derived mAb, MuSK3B, was not further investigated in this study, as this isotype has not been implicated in MuSK MG pathology. Given the importance of IgG4 autoantibodies in MuSK MG and to discount any influence of the subclass constant region on the IgG4 mAbs, we sub-cloned the variable region of the IgG4-subclass autoantibodies (MuSK1A and MuSK3-28) into human IgG4 expression vectors. Thus, matching the native subclass and ensuring they were expressed as divalent, monospecific recombinant mAbs. Unless otherwise noted, the IgG4-subclass versions of mAbs MuSK1A and MuSK3-28 were used.

Binding of the mAbs was tested over a range of concentrations using a live CBA. These tests demonstrated that binding could be detected with only 20 ng/mL for the IgG mAbs MuSK1A, MuSK1B and MuSK3-28 **(Figure 2A, B**). An independent radio-immunoprecipitation assay (RIA), commonly used for clinical diagnosis of MuSK MG, showed that as little as 0.3 ng of mAbs MuSK1A, MuSK1B and MuSK3-28 could bind 30-50% of ^125^I-MuSK (approximately 1 fmol/assay) **(Figure 2C)**. To explore their specificity and potential pathogenicity, we also tested these mAbs in CBAs using GFP-transfected HEK cells, or cells transfected with AChR or MOG **(Figure 2D)**. MOG was chosen because its structure (50), a classical Ig (Ig variable domain) fold, is highly similar to that of MuSK (51). The mAbs MuSK1A, MuSK1B, and MuSK3-28 did not show any detectable binding to GFP-transfected HEK cells or HEK cells expressing AChR or MOG on their surface **(Figure 2D)**. Finally, we tested the mAbs on sections of mouse muscle tissue to determine whether they could recognize mouse MuSK. The mAbs that were highly positive in the CBA, using human MuSK, also bound to mouse NMJs where the mAbs closely colocalized with AChRs **(Figure 2E).** These data also demonstrated that the mAbs can recognize MuSK when presented in its native biological environment, an important requisite for future pathogenicity experiments in mouse-derived cells and disease models.

**Figure 2.**
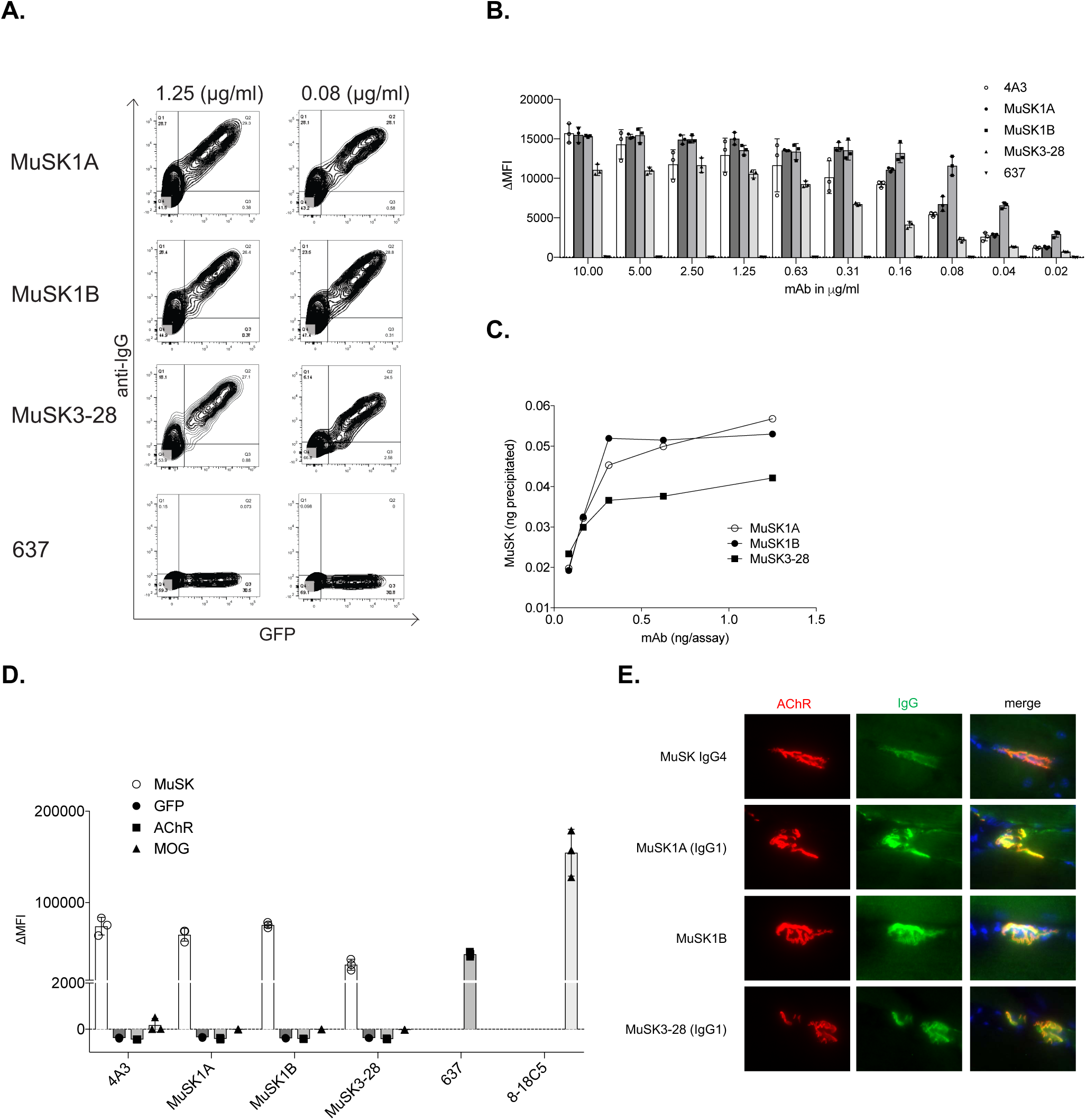
Characterization of human MuSK mAb binding properties. Binding properties of mAbs MuSK1A, MuSK1B and MuSK3-28 were tested in several *in vitro* antibody binding assays. **A**. Representative cell-based assay (CBA) flow cytometry plots are shown for three MuSK mAbs and a negative control (AChR-specific mAb 637). Binding was tested at both 1.25 and 0.08 μg/mL. The x-axis represents GFP fluorescence intensity and, consequently, the fraction of transfected HEK cells. The y-axis represents Alexa Fluor 647 fluorescence intensity, which corresponds to secondary anti-human IgG antibody binding and, consequently, primary antibody binding to MuSK. Hence, transfected cells are located in the right quadrants and transfected cells with MuSK autoantibody binding in the upper right quadrant. **B.** Binding to MuSK was tested over a wide range of mAb concentrations in the CBA. Controls included the MuSK-specific humanized mAb 4A3 and AChR-specific mAb 637 tested with MuSK mAbs MuSK1A, MuSK1B and MuSK3-28. Each data point represents a separate replicate within the same experiment. Bars represent means and error bars SDs. **C.** A solution phase radioimmunoassay (RIA) was used to measure MuSK binding over a range of mAb concentrations. Each data point represents a value within the same experiment. **D.** Specificity of the mAbs was evaluated using CBAs that tested binding to HEK cells transfected with MuSK, GFP alone, AChR or myelin oligodendrocyte glycoprotein (MOG). Positive controls included MuSK-specific humanized mAb 4A3, AChR-specific mAb 637, and MOG-specific 8-18C5. Each data point represents a separate replicate within the same experiment. Bars represent means and error bars SDs. **E.** Immunofluorescent staining of mouse neuromuscular junctions. Tibialis anterior muscles were cut longitudinally in cryosections and fixed with PFA. AChRs were stained with Alexa648-alpha-bungarotoxin (shown in red), and DNA with Hoechst (shown in blue in the merged panels). The first row shows staining with polyclonal IgG4 from a MuSK MG patient. Binding of mAbs (MuSK1A, MuSK1B, MuSK3-28) against MuSK (1.6 µg/mL for 1 hr) was detected with goat anti-human IgG Alexa488 (mAb, shown in green). In panels **A-E** the IgG4 subclass mAbs MuSK1A and MuSK3-28 were tested in their native IgG subclass unless indicated otherwise.

### MuSK autoantibody epitope mapping

The extracellular domain of MuSK is comprised of three Ig-like domains (Ig1, 2 and 3) and a cysteine-rich frizzled-like domain (Fz-like), which occupies the region between the Ig-like domains and the extracellular juxtamembrane region (51, 52). The majority of serum-derived MuSK autoantibodies are reported to recognize the N-terminal Ig-like domain-1, which interacts with LRP4 (20). The interruption of Agrin-induced clustering of the AChR by MuSK autoantibodies is likely to be mediated by autoantibodies that recognize this domain. However, serum-derived autoantibodies that recognize other domains of MuSK have been reported (7, 20, 53). To map the targets of the mAbs, we engineered a series of plasmid constructs (**Figure 3A and Figure S3**) to express either individual subdomains of MuSK or MuSK with deletions of individual subdomains and tested binding of the mAbs using a CBA. The mAbs MuSK1A, MuSK1B and MuSK3-28 bound to HEK cells expressing full-length MuSK, MuSK ΔIg-1, MuSK ΔFz, and MuSK Ig2 only (ΔIg1, 3 and Fz) (**Figure 3B**), but did not bind to MuSK in which the Ig-like domain 2 was deleted (ΔIg2) or when the Ig-like domain 1/Fz-like domain were expressed alone (**Figure 3B**). These data indicate that mAbs MuSK1A, MuSK1B and MuSK3-28 recognize epitope(s) in Ig-like domain 2. The control humanized mAb 4A3, which we produced (32), recognized the Fz-like domain (**Figure 3B**).

**Figure 3.**
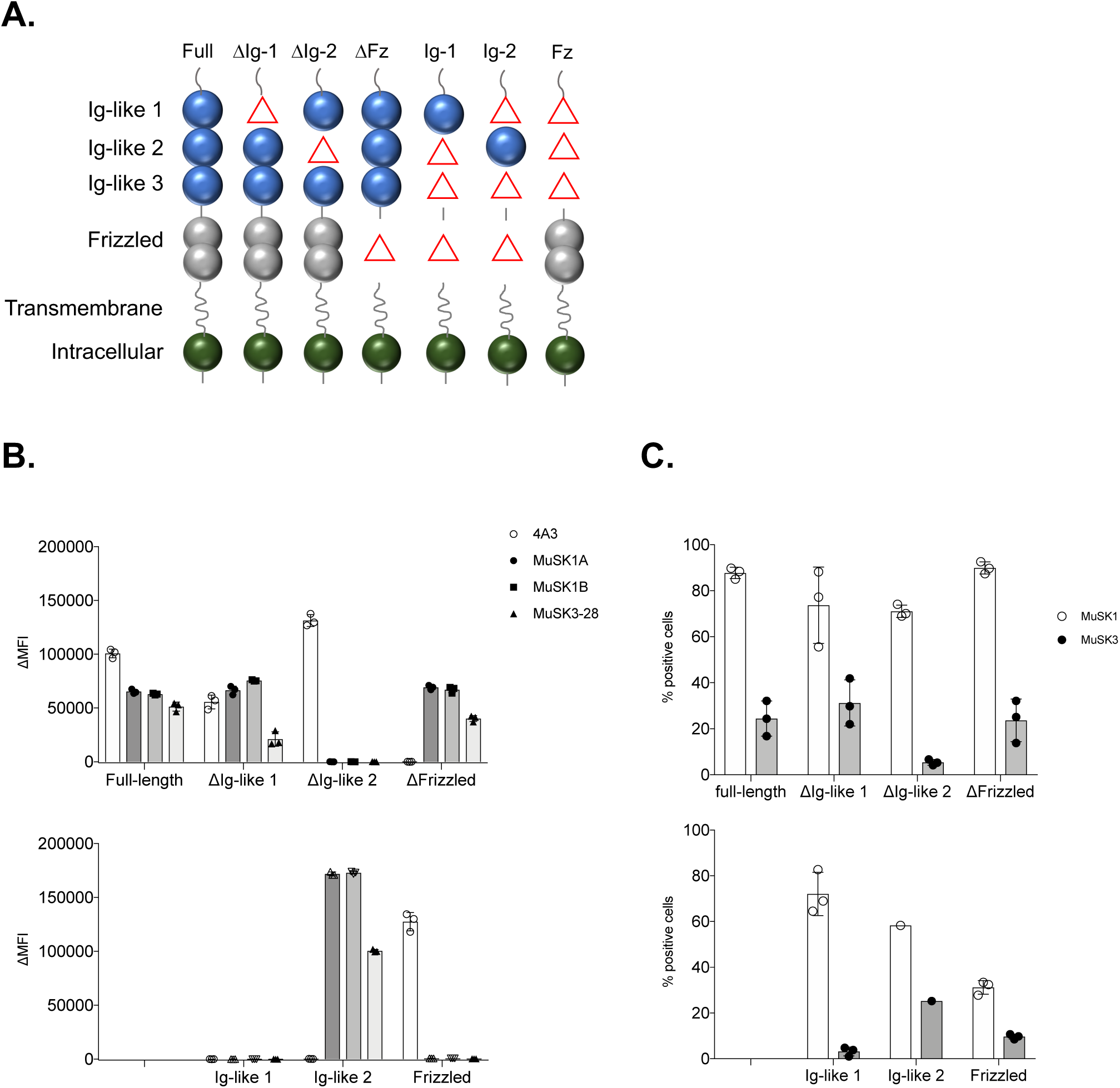
MuSK domain binding results. To map the human MuSK mAb epitopes, MuSK constructs that had particular domains deleted and full-length MuSK were each expressed in HEK cells and tested with the cell-based assay (CBA). The schematic in (**A**) illustrates the mutant forms of MuSK. For example, “ΔIg-1” includes only the Ig-like domains 2 and 3, and the frizzled-like domain, as the Ig-like domain 1 was deleted (shown as a “Δ” in the schematic). Similarly, “Ig-1” includes only the Ig-like domain 1 as the Ig-like domains 2, 3 and frizzled-like were deleted (shown as “Δ” in the schematic). Binding of mAbs (MuSK1A, MuSK1B, MuSK3-28, and the positive control humanized MuSK mAb 4A3) to these mutant forms of MuSK was tested in our standardized flow cytometry CBA. Results for each mAb (**B**) or serum **(C)** specimen are shown. Serum was obtained from the same patients from whom the mAbs were derived. Each data point represents a separate replicate within the same experiment. Bars represent means and error bars SDs.

We also tested sera from MuSK MG patients MuSK1 and MuSK3 for binding to the different MuSK domain constructs. MuSK1 serum contained autoantibodies that recognized full-length MuSK, as well as each of the domain deletion constructs and the isolated domain constructs. These findings indicate that serum contains a heterogeneous collection of autoantibody specificities, which collectively recognize epitopes present in all of the tested MuSK domains, (**Figure 3C**). MuSK3 serum displayed lower reactivity compared to MuSK1 serum when testing binding to full-length MuSK. In addition, MuSK3 serum contained autoantibodies that preferentially recognize MuSK mutants that included the Ig-like domain 2, indicating that the epitope(s) could be more restricted in this patient (**Figure 3C**). These results using either MuSK1 and MuSK3 serum provide further evidence for surface expression of the constructs tested. Moreover, the MuSK3 serum binding results showed that the specificity of the mAb MuSK3-28 for the MuSK Ig-like domain 2 was reflected in the circulating autoantibody repertoire.

### Pathogenic capacity - MuSK mAbs interfere with Agrin-induced AChR clustering

To evaluate the pathogenicity of the MuSK-specific recombinant mAbs we used the well-established *in vitro* C2C12 AChR clustering assay. The C2C12 mouse myotubes express all of the components that are required for Agrin to stimulate AChR clustering (54), and is dependent on functional MuSK (*MC, AV and DB unpublished data*). Serum-derived MuSK autoantibodies have been demonstrated to interrupt this interaction and consequentially inhibit AChR clustering (7, 18, 23).

The C2C12 myoblasts were differentiated into myotubes and incubated with each of the three MuSK mAbs and controls in duplicate wells. AChR clusters were visualized (**Figure 4A-D**) using Alexa Fluor 647-conjugated α-bungarotoxin and counted; the mean number of clusters in four 100X magnification visual fields was recorded. These data (**Figure 4E**) show that all three mAbs reduced Agrin-induced AChR clustering, whereas mAb 4A3, which recognized the Fz-like domain, had no effect. These findings indicate that mAbs MuSK1A, MuSK1B and MuSK3-28 are pathogenic in this model.

**Figure 4.**
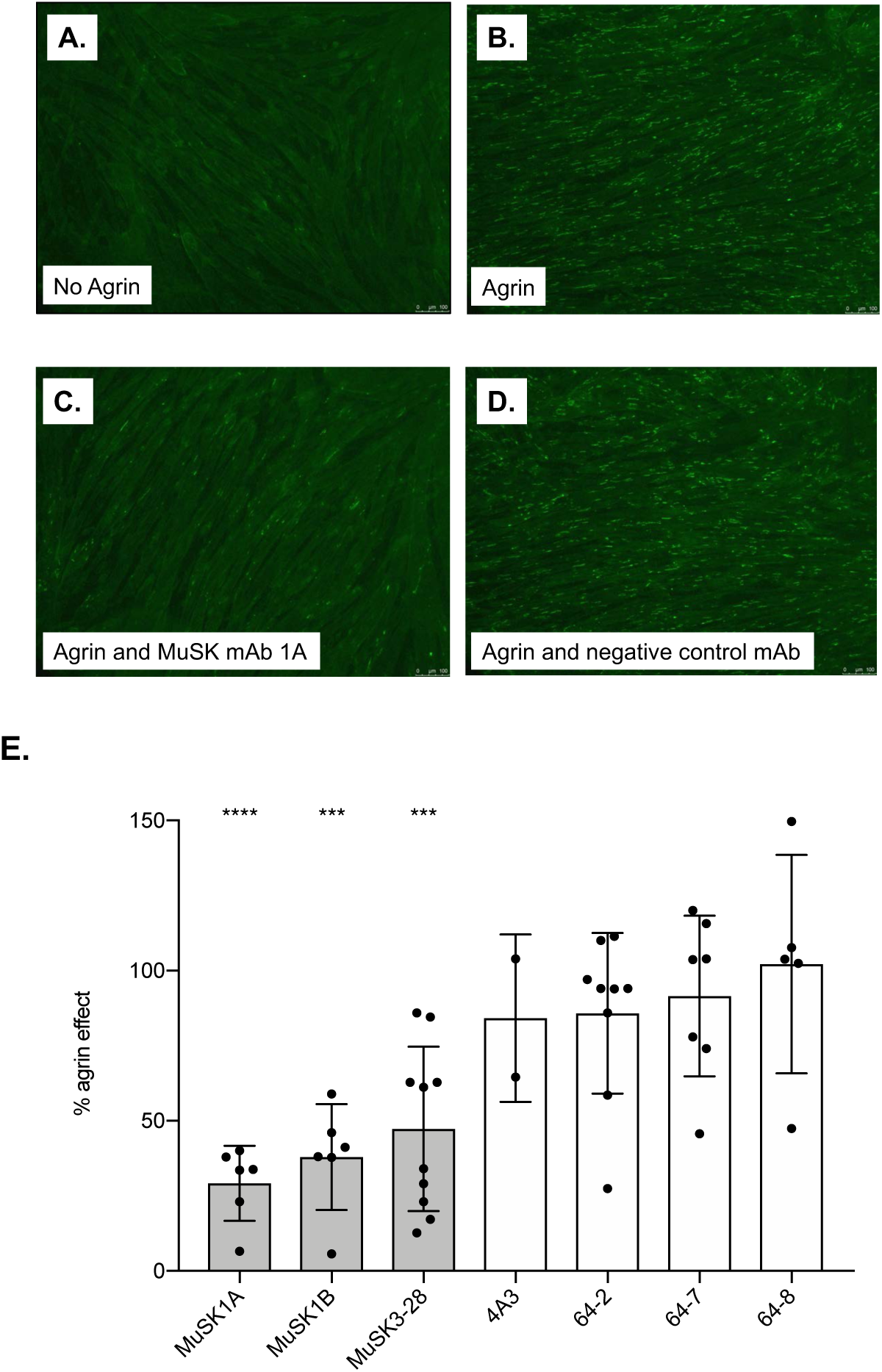
AChR clustering assay in C2C12 mouse myotubes demonstrates pathogenic capacity of MuSK mAbs. The presence of agrin in C2C12 myotube cultures leads to dense clustering of AChRs that can be readily visualized with fluorescent α-bungarotoxin and quantified. Pathogenic MuSK autoantibodies disrupt this clustering. Three different human MuSK-specific mAbs, the humanized murine control MuSK mAb 4A3 and three human non-MuSK-specific mAbs derived from AChR MG patient plasmablasts (64-2, 64-7 and 64-8) were tested for their ability to disrupt the AChR clustering. Each mAb was added to the cultures at 1μg/mL. Representative images from the clustering experiments are shown (**A-D**). Cultured myotubes (**A**) do not show AChR clustering until agrin (**B)** is added (*bright spots reveal AChR clusters*). The mAb MuSK1A added at 1μg/mL inhibits clustering (**C**), while a control mAb does not inhibit the formation of AChR clusters (**D**). Clustering of AChR was quantified relative to the measured effect of agrin. Quantitative results are normalized to agrin-only induced clustering (**E**). Each data point represents the mean value from an independent experiment. Bars represent the mean of means and error bars SDs. Multiple comparisons ANOVA (against the pooled results for the three human non-MuSK-specific mAbs), Dunnet’s test; * p<0.05, ** p<0.01, *** p<0.001, **** p<0.0001, only shown when significant.

The ability of the mAbs to induce AChR clustering in the absence of Agrin was also evaluated. When added to C2C12 cultures, each of the three MuSK mAbs induced a modest increase in AChR clustering **(Figure S4)** compared to three non-MuSK binding mAbs and the MuSK Fz-like domain-specific 4A3 mAb.

### Pathogenic capacity - MuSK mAbs modify Agrin-induced MuSK phosphorylation

One of the crucial steps in the Agrin/LRP4/MuSK/DOK7 pathway is MuSK phosphorylation. Serum IgG or IgG4 antibodies from patients with MuSK MG inhibit Agrin-induced MuSK tyrosine phosphorylation (20). MuSK MG serum-derived IgG or recombinant human mAbs were added to cultured C2C12 myotubes, together with Agrin. MuSK tyrosine phosphorylation was then detected using immunoblotting with a phosphotyrosine-specific antibody (**Figure 5A)**. Agrin-induced phosphorylation was blocked by the patient-derived serum IgG whereas the three non-MuSK binding mAbs and the control mAb 4A3, which recognizes an epitope in the Fz-like domain, did not alter the Agrin-induced MuSK phosphorylation (**Figure 5B**). By contrast, and intriguingly, the three MG patient-derived MuSK specific mAbs, MuSK1A, MuSK1B and MuSK3-28, all modestly amplified the Agrin-induced phosphorylation (**Figure 5B**). Thus, these mAbs increased Agrin-induced MuSK phosphorylation while inhibiting Agrin-induced AChR clustering. The mAbs MuSK1A and MuSK3-28 were tested both as their native IgG4 subclass and also as IgG1. Similar amplification of Agrin-induced phosphorylation was observed with both subclasses (**Figure 5B)**. These results suggest that divalent (and monospecific) MuSK autoantibodies that bind Ig-like domain 2 can activate MuSK phosphorylation, irrespective of their subclass.

**Figure 5.**
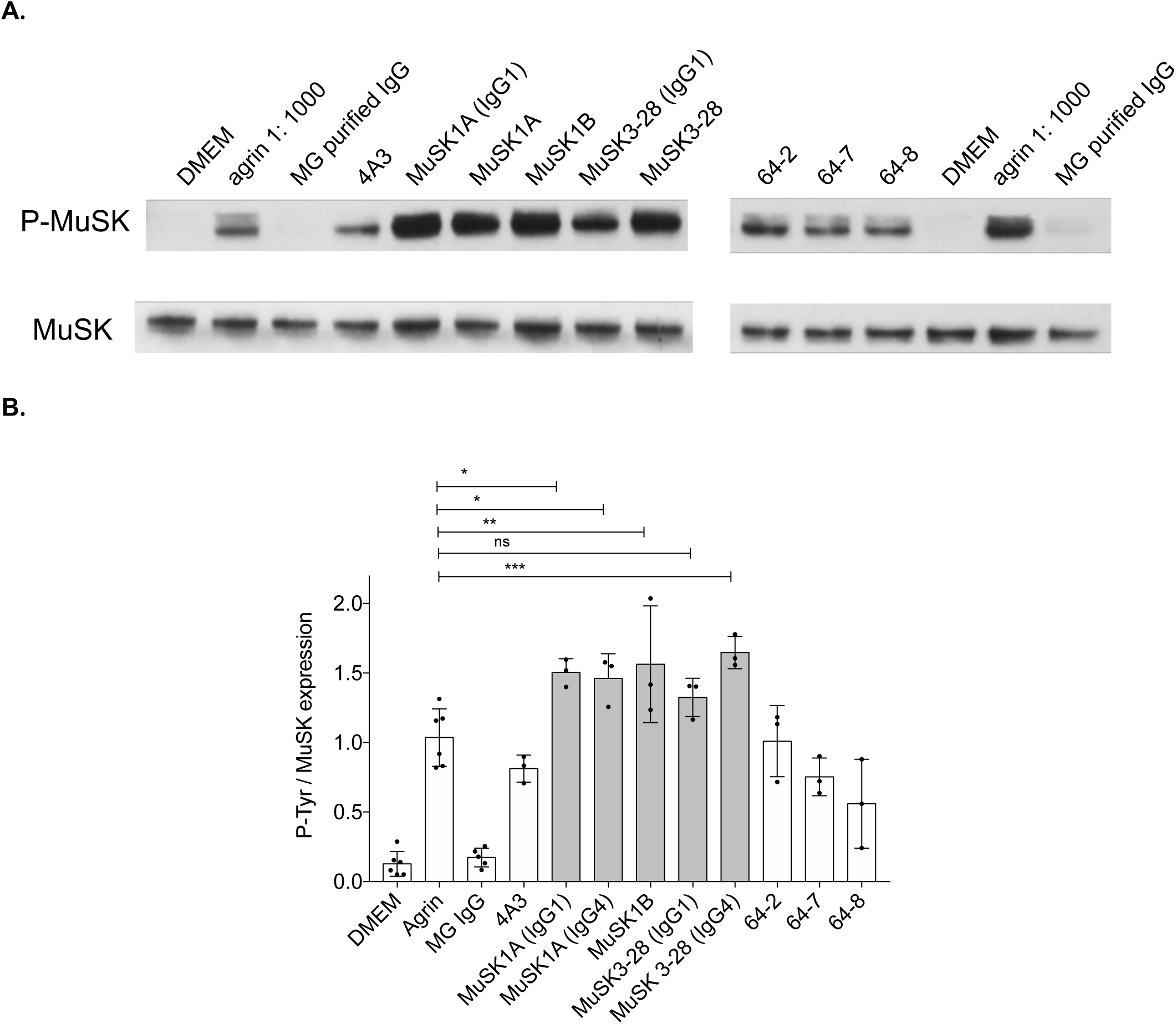
MuSK mAbs can amplify agrin-induced tyrosine phosphorylation. A. Immunoblots showing phospho-tyrosine bands and related MuSK expression in C2C12 murine myotubes that were incubated with agrin in presence of MuSK MG serum-derived IgG4 or recombinant MuSK/control mAbs. 4A3 is a humanized murine MuSK mAb, MuSK1A, MuSK1B, and MuSK3-28 are human MuSK mAbs from MuSK MG patients, and 64-2, 64-7, and 64-8 are non-MuSK binding human mAbs derived from AChR MG patient plasmablasts. IgG4 subclass mAbs MuSK1A and MuSK3-28 were expressed in vectors reflecting the native subclass and also as IgG1 (as indicated). **B**. Normalized densitometry analysis results from the MuSK phosphorylation immunoblots are plotted. Each data point represents an independent experiment. Bars represent means and error bars SDs. Phosphorylation of MuSK was determined by normalizing to MuSK expression, detected by a commercial anti-MuSK antibody after stripping the blot, and the ratio of phosphor-tyrosine MuSK to MuSK is plotted. Multiple comparisons ANOVA (versus agrin), Dunnet’s test; ns p>0.05, * p<0.05, ** p<0.01, *** p<0.001, shown for MuSK mAbs vs. agrin comparisons.

## Discussion

The value of isolated human autoantibody-producing B cells in advancing the understanding of immunopathology is considerable. Individual autoantibody-producing cells and production of recombinant human monoclonal autoantibodies from these cells allow questions, which were previously unapproachable, to be explored. This is highlighted by the cloning of unique AChR autoantibody-producing B cells, which was essential to gain understanding of AChR MG (37, 55-59) disease mechanisms. Similarly, features of neuromyelitis optica immunopathology have been replicated in animal models through the transfer of human-derived autoantibody mAbs, demonstrating the direct role these autoantibodies play in the disease (60). However, the identification and isolation of autoantibody-producing cells is challenging because many reside in the lymphatics, bone marrow or other tissue compartments and are often scarce in the circulation (33, 61). Nonetheless, circulating antigen-specific B cells can be detected in autoantibody-mediated conditions (62, 63). Screening approaches for these cells, such as using Epstein-Barr virus transformation of circulating B cells or bulk isolation of antigen-experienced B cells, provide low yields of desired autoantibody specificities (64-66), thus illustrating the difficulty in the isolation of such cells. Here, we used a MuSK construct that included a BirA tag, which enabled post-expression multimerization and fluorescent labeling, to identify, capture, and investigate the functional relevance of the autoantibody-producing B cells. Our approach, while enriching for autoantibody-producing B cells, also highlighted the rarity of such cells in the circulation. The low yield notwithstanding, we identified autoantibodies that in CBAs demonstrated binding to MuSK with ΔMFIs that were 250-fold higher than HD-derived mAbs at 1 μg/mL and in RIAs bound to MuSK at sub-nanomolar concentrations. Further investigations, which were outside of the scope of this study, will be required to understand the relevance of the many low-strength binders we isolated from patients and HDs.

Our characterization of several IgG MuSK mAbs focused on their binding properties, molecular properties and pathogenic potential. Our exploration of these characteristics revealed that MuSK MG immunopathology includes mechanistic details that were not readily appreciated through the study of polyclonal immunoglobulin derived from human serum of MuSK MG patients. Each of these mAbs bound MuSK expressed in live transfected HEK cells, which preserves biological integrity and native antigenicity of the membrane proteins and is used extensively in diagnostic assays for many autoantibody-mediated diseases targeting membrane receptors or ion channels (67). We unambiguously authenticated the binding in independent and complementary assay platforms. In contrast to the predominant epitope specificity (Ig-like 1 domain) reported elsewhere, the three IgG mAbs specifically recognized the Ig-like 2 domain of MuSK, while the humanized murine control mAb 4A3 recognized the Fz-like juxta-membrane domain. Nevertheless, the three IgG mAbs inhibited the formation of Agrin-induced AChR clusters, in the C2C12 myotube model and bound to MuSK at the mouse NMJs, suggesting that they are likely to have full pathogenic potential and perhaps unique downstream molecular mechanisms. These novel findings raise questions regarding the pathogenic dominance of monovalent IgG4 autoantibodies to the Ig-like 1 domain in MuSK MG.

Overall, this study shows that isolated human monoclonal autoantibodies, which bind MuSK, are sufficient to perturb Agrin-induced clustering of the AChR, consistent with the well-accepted mechanism of MuSK MG. Consequently, it provides evidence that MuSK autoantibodies alone are capable of affecting the immunopathology and that it does not depend on the presence of additional serum proteins such as complement factors or co-existing autoantibodies with other specificities (68-70).

### MuSK autoantibody-producing B cell phenotypes

Four mAbs that bound to MuSK in a robust manner were successfully isolated from two unique patients, while three additional patient samples did not yield any strong MuSK-binding mAbs. These results point toward the remarkable rarity of such cells in the circulation of patients, even though such patients were experiencing active disease and had conspicuous serum autoantibody titers at the time of specimen collection. The mAb MuSK1B was derived from a cell with a plasmablast-like phenotype, expressing both CD27 and high levels of CD38. This plasmablast, along with that which produced mAb MuSK3-28 (32), support the notion that this circulating short-lived cell-type contributes to MuSK MG immunopathology. We previously proposed (31) that these cells are either direct or indirect targets of CD20-mediated B cell depletion depending on whether they express CD20 or are continuously supplied by CD20-bearing cells.

The isolated cell which yielded mAb MuSK1A displayed a memory B cell-like phenotype. The finding of this cell type among MuSK autoantibody-producers suggests that immunological memory might have been established in the MuSK MG patient it was derived from. One can speculate that activated memory B cells, such as that which produced mAb MuSK1A, can provide a reservoir from which autoantibody-secreting plasmablasts originate. Given that memory B cells express CD20, their direct elimination by CD20-targeted therapy (rituximab) may be the mechanism by which this treatment effects remarkable serum autoantibody decline and excellent clinical response in patients.

The patient from whom this B cell was isolated had received rituximab, responded well, then relapsed and it was during this relapse that the sample was collected. It will be interesting to learn whether the memory B cell, producing mAb MuSK1A, was historic or recently generated. Antibody repertoire sequencing, which accurately identifies B cell clones through their unique B cell receptor sequence, could provide such information through analysis of specimens collected prior to treatment, but these data were not available for this study. The identification of historical clones would indicate that they escape B cell depletion, then contribute to clinical relapse after the treatment is withdrawn. Evidence supporting this mechanism has been recently reported: Tissue-resident vaccine-specific memory B cells, which are refractory to depletion, undergo re-vaccination-induced expansion during B cell repopulation following rituximab treatment (71). On the other hand, *de novo* generation may be occurring which would point toward a fundamental defect in B cells that, once generated, can efficiently re-establish autoantibody production and consequently disease manifestation.

### MuSK autoantibody IgG subclasses and epitope recognition

The majority of MG patients have autoantibodies against the AChR, which belong predominantly to the IgG1 and IgG3 subclasses. These autoantibodies activate the complement system and can cross-link the AChR resulting in endocytosis. In contrast, serum MuSK autoantibodies are reported to be predominantly IgG4 (14, 72); this IgG subclass neither activates complement, nor efficiently cross-links antigens (15, 23). The first Ig-like domain in MuSK, is required for MuSK to bind LRP4 (73). MuSK IgG4 serum autoantibodies cause MG by inhibiting binding between MuSK and LRP4 (20, 21). Other IgG subclasses, including IgG1-3, although far more rare than the IgG4 subclass, can inhibit Agrin-induced clustering in C2C12 myotubes (21). The inability to cross-link MuSK arises because IgG4 molecules exchange Fab arms with other IgG4s, thus the products are effectively functionally monovalent. In the Fab-arm exchange reaction a monospecific IgG4 immunoglobulin may swap a heavy and light chain pair with another IgG4 to become bispecific (24). Fab-arm-exchanged antibodies are present in MuSK MG patient serum and these antibodies reduced AChR clustering in the C2C12 model (23), as do polyclonal Fabs against MuSK (21). Because our IgG4 mAbs were expressed as individual clones, they are not expected to participate in the Fab-arm exchange reaction, and are thus divalent (and monospecific). We found that two autoantibodies (MuSK1A and MuSK3-28) are of the IgG4 subclass; as would be expected considering the results of many studies of MuSK MG serum (14, 72). However, one (MuSK1B) is from the IgG3 subclass. These findings present the opportunity to widen our understanding of MuSK autoantibody-mediated pathology. In their divalent (and monospecific) form, which we tested here, the IgG4 (expressed with an IgG4 constant region) and IgG3 (expressed with an IgG1 constant region) subclass mAbs all effectively inhibited Agrin-induced AChR clustering. These data suggest that all have pathogenic potential. Interestingly, the same mAbs amplified MuSK phosphorylation, while MuSK MG serum IgG effectively blocked Agrin-induced MuSK phosphorylation. Thus, these mAbs behave in a manner opposite to that which is observed with serum-derived IgG. Given their recombinant expression, all of our mAbs were divalent and monospecific and consequently they effectively cross-linked MuSK, which has been shown to affect phosphorylation in murine models (19, 74, 75). The IgG4 mAbs may participate in Fab-arm exchange (predominantly with IgG4 antibodies of other specificities) *in vivo* and thus could still inhibit clustering but not cross-link MuSK and thereby not amplify autophosphorylation. The IgG3 mAb (MuSK1B) would not be expected to participate in Fab-arm exchange, but instead it could activate the complement system at the NMJ if it bound there at sufficient density to allow C1q binding. While it may interrupt AChR clustering through steric hindrance of LRP4-MuSK interaction, the contribution of the agonistic effect, observed in the phosphorylation experiments, to pathology is not clear. The use of *in vivo* models (discussed below) will likely be required to unravel these roles.

The N-terminal Ig-like domain of MuSK is crucial for the interaction with LRP4 and ultimately for AChR clustering and NMJ maintenance (53). It follows that the proposed pathogenic mechanism of serum-derived MuSK autoantibodies interferes with this process through specific binding of the N-terminal Ig-like domain (22). Indeed, serum from patients with MuSK MG most often and more robustly bind to the N-terminal Ig-like domain, but some binding to other regions of the molecule has been observed (14, 53). We found that the three MuSK mAbs (MuSK1A, MuSK1B and MuSK3-28) recognize the Ig-like domain 2, not the more commonly recognized Ig-like 1 domain, while the humanized murine control MuSK mAb 4A3 recognizes the Fz-like domain. It has not been determined whether autoantibodies that recognize Ig-like domain 2 mediate pathology. It is possible that binding to the Ig-like domain 2 also hinders MuSK-LRP4 binding.

The amplified Agrin-induced phosphorylation may be associated with the binding to the Ig-like domain 2. We reason the mAbs dimerized MuSK, by nature of their bivalency and monospecificity, which led to the autophosphorylation, as this has been observed through using agonistic MuSK mAbs (19, 74, 75). However, it remains to be determined how binding to domains outside of Ig-like domain 1 mediated downstream activation of MuSK and other molecules in the NMJ.

### Limitations

Several important limitations require consideration. The first involves pathogenic contributions. We have isolated B cells and plasmablasts that express MuSK autoantibodies, however the contribution they make to serum autoantibodies in the patients has not been established. Empirical evidence demonstrating that individual B cells or plasmablasts contribute to the reservoir of pathogenic autoantibodies in the serum is required to establish that these cells effect the disease process. This can be achieved through leveraging the unique antibody sequences that identify a B cell clone. These sequences can be compared to those derived from a proteomic analysis of the serum autoantibody repertoire within the same patient. This approach has been performed in several studies which have shown close associations between the B cell and the immunoglobulin repertoire (76) and others that point toward more diversity in the serum immunoglobulins (77). Three of the MuSK mAbs demonstrated pathogenic capacity in the AChR clustering assay. The same mAbs show agonistic activity in the MuSK phosphorylation assays. These findings using *in vitro* cellular models indicate that the mAbs are functionally active, but further study is required to more fully understand the mechanisms by which they might contribute to disease *in vivo*. Pathogenicity can be evaluated *in vivo* using well-established passive transfer models of MuSK MG (16, 17). Not only will such *in vivo* studies further define the pathogenic capacity of the mAbs, they will provide additional insights, such as the effects on particular muscle groups. Finally, since IgG4 antibodies can be functionally monovalent as a result of Fab-arm exchange (23, 24), experiments assessing pathogenic capacities of MuSK1A and MuSK3-28 *in vitro* and *in vivo* under functionally monovalent conditions will be required to more thoroughly understand how they impact immunopathology.

### Broader implications of impact

We anticipate that these human MuSK mAbs and the approach to their isolation will be recognized as highly valuable tools in future studies. First, these mAbs can be used to dissect the molecular mechanisms of MuSK autoantibody pathology through studying how they specifically interact with MuSK and influence changes at the NMJ. Such detailed molecular dissection of autoantibody structure can potentially provide information for the design of treatments (such as molecular decoys or cell-based therapies) that target the autoimmune response with precision. Second, novel, sustainable pre-clinical models can be developed that do not rely on rare and limited human MG-derived serum autoantibodies. Specifically, clinically relevant animal models are expected to be developed, based on the passive transfer of defined, MuSK mAbs that are identical to some of the patient autoantibodies and will be available in unlimited supply at constant quality. Third, the identification and isolation of rare MuSK mAb-producing cells, using the fluorescent MuSK antigen, will allow further investigation into their cellular dysfunction. Of particular interest is understanding the activation state of these autoreactive B cells through examination of their gene expression profile, how they escape tolerance mechanisms and why they generally fail to establish a long-lived plasma cell compartment. In addition, their frequency in the circulation may represent a valuable biomarker for predicting relapse and therapeutic response. Other IgG4-mediated autoimmune diseases are thought to utilize similar cellular mechanisms of immunopathology, such as pemphigus and chronic inflammatory demyelinating polyneuropathy (CIDP) (78, 79). Thus, these MuSK autoantibody-producing B cells may provide insight into similar autoimmune pathologies. Finally, these cells are viable targets of personalized medicine that would seek to eliminate only those cells that directly contribute to autoimmunity. The obvious advantage of which is to circumvent collateral damage to B cell populations that are needed for established immunity and are targeted by current immune modulating treatments.

## Acknowledgments

The authors thank Professor Steven J. Burden and Dr. Damian Ekiert of the Skirball Institute of Biomolecular Medicine, New York University Medical School, for critically reading the manuscript.

## Author contributions

This study was originally conceived, then initiated and directed by KCO. RJN provided all characterized subject specimens and directed the clinical aspects of the study. KT produced the tetramer. KT and PS performed the single cell isolation, mAb expression, mAb sequencing and cell-based antibody assays and interpreted those data. MC, DB and AV designed, performed, and interpreted the phosphorylation assays. MMD, PMM and ML designed and performed the murine muscle staining and interpreted those data. LJ, PW, SRI and AV performed the radioimmunoassays and interpreted those data. MF performed additional cell-based antibody assays and contributed to the C2C12 assays that were established and performed by PS. KT, with assistance from EB, built and tested the MuSK domain deletion constructs. Data were analyzed and interpreted by all authors. All authors contributed to manuscript editing after it was initially drafted by KCO and AV with input from PS and ML. Key contributions in editing were further provided by PS and ML.

## Supplemental data to accompany manuscript

**Figure S1.**
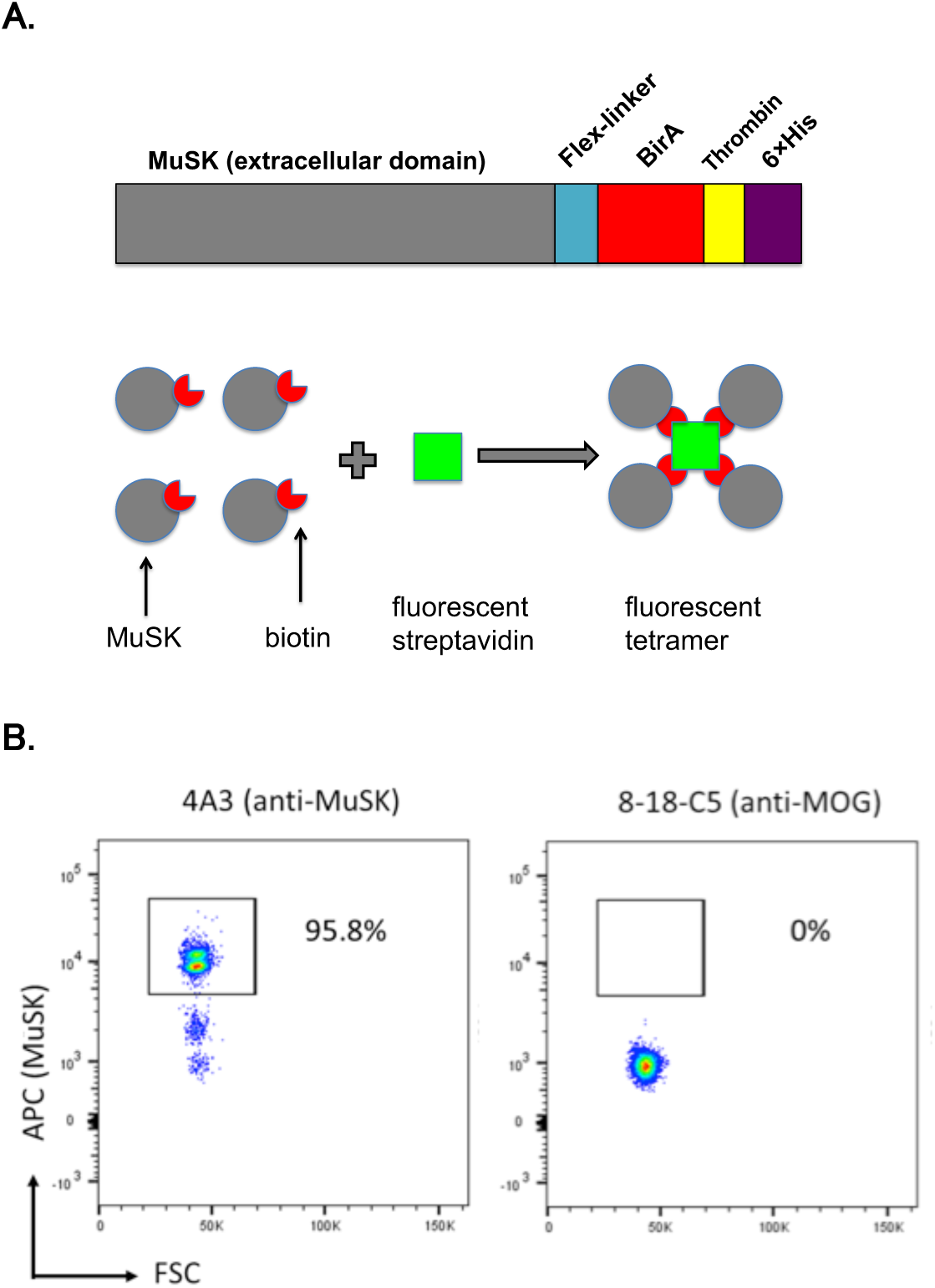
Multimerized, labeled MuSK retains properties required for antibody recognition and detection by flow cytometry. Schematic diagram **(A)** showing MuSK expression construct and post-translational tetramer assembly. FACS plots **(B)** showing feasibility of MuSK tetramer-mediated isolation. Flow cytometry beads coated with murine monoclonal antibodies that recognize MuSK (4A3) or myelin oligodendrocyte glycoprotein MOG (8-18C5) were stained with APC-conjugated MuSK multimer. Detection of binding to the beads was analyzed by flow cytometry. Data shown is one of two representative experiments.

**Figure S2.**
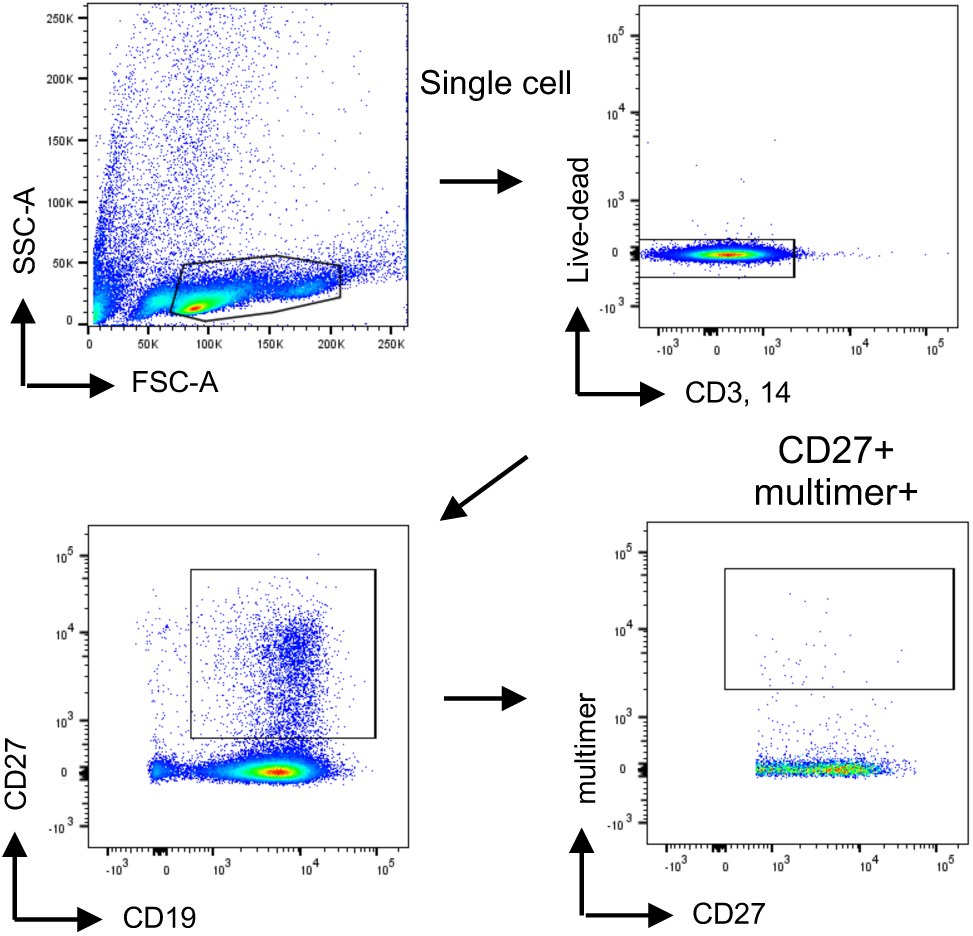
MuSK multimer FACS gating strategy. A representative example of the MuSK multimer positive population sorting strategy is shown. B cells that were enriched through bead-based negative selection, were initially gated in the SSC/FSC graph. After doublets and dead cells were excluded, CD19+CD3-CD14- cells were gated. MuSK multimer positive cells were subsequently gated from the B cell gate (CD19+CD3-CD14-) using CD27+ MuSK multimer+ cells. Plasmablasts were defined as CD27hiCD38hi cells for purposes of back-gating and index-sorting (not shown).

**Figure S3.**
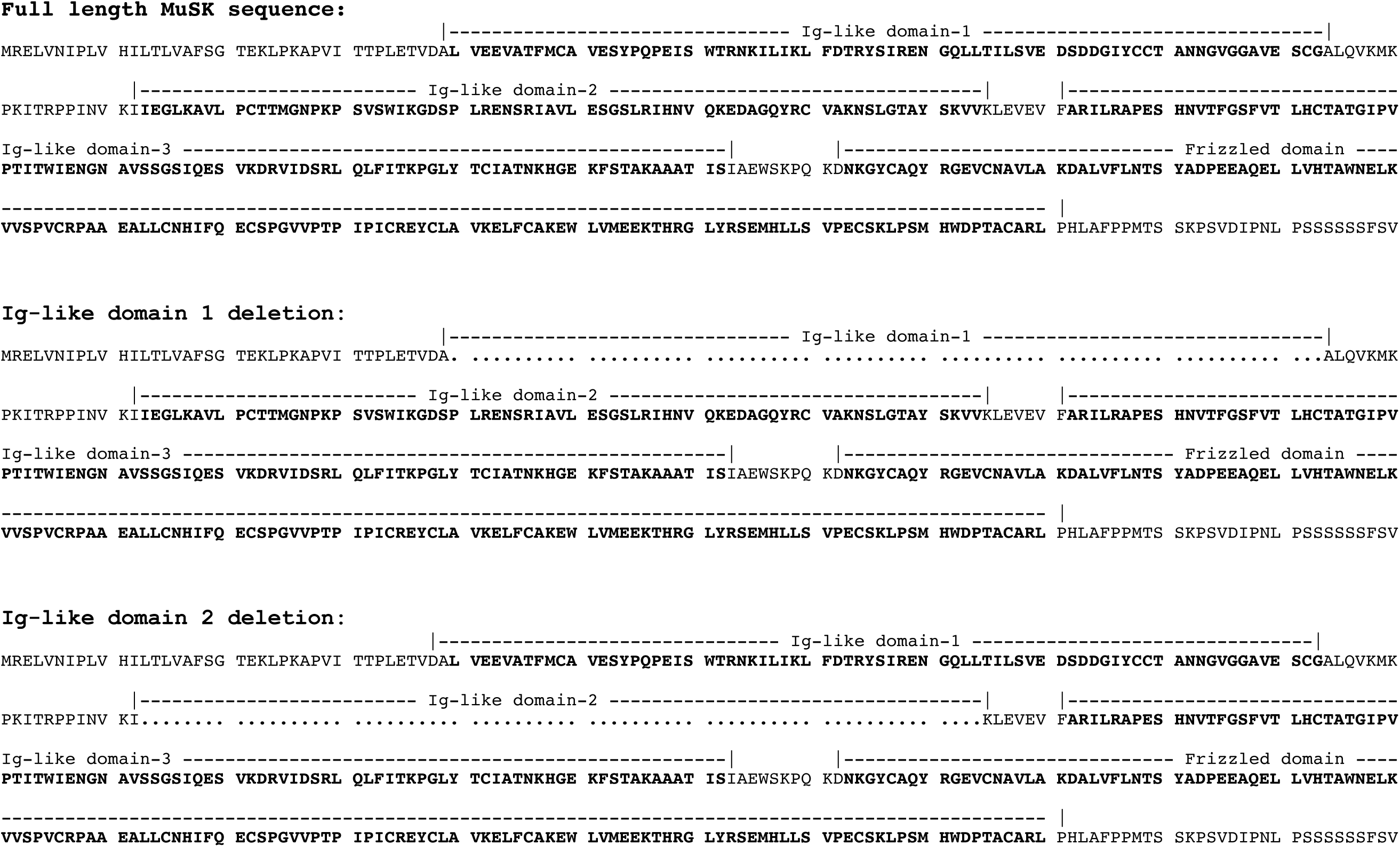
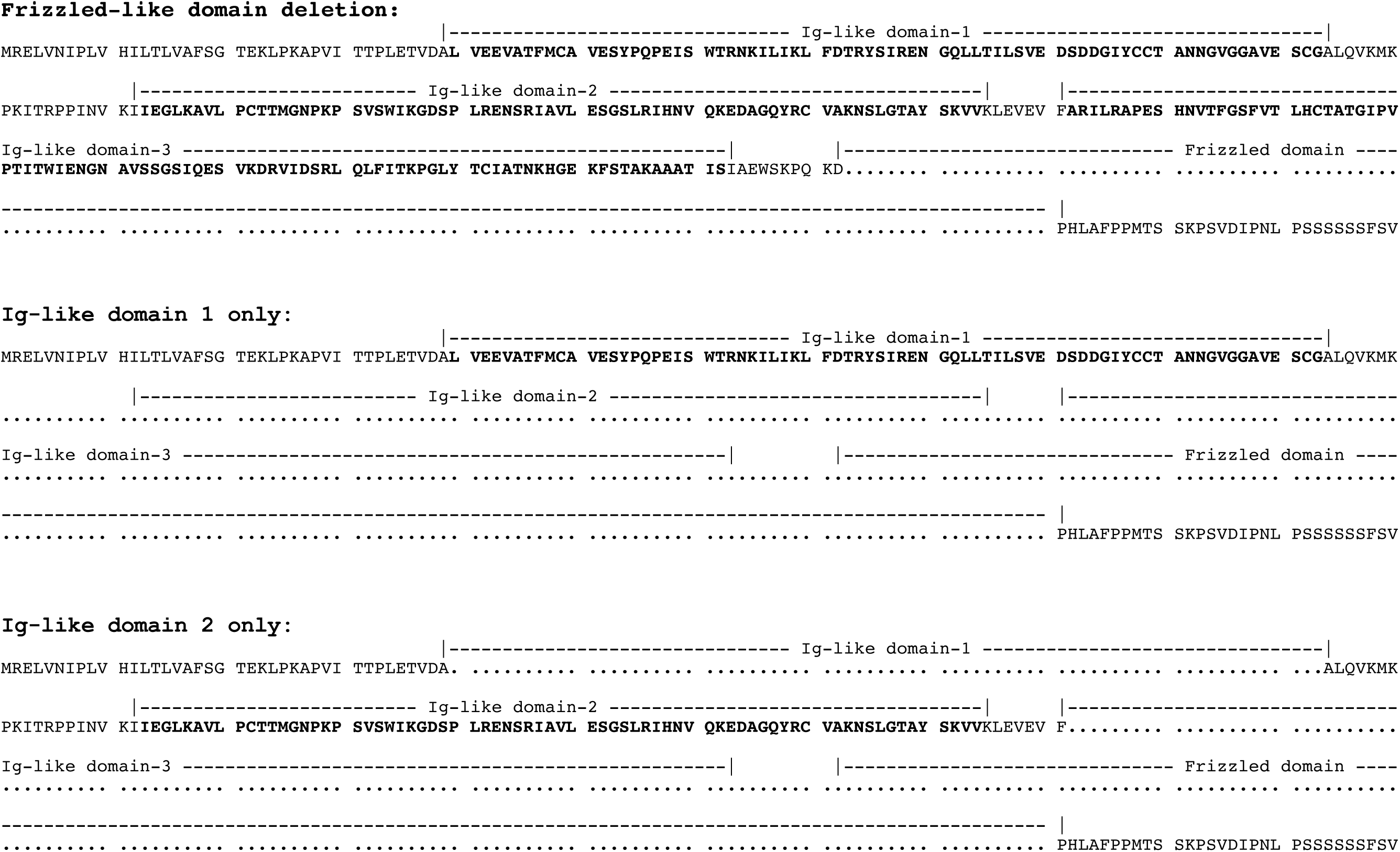
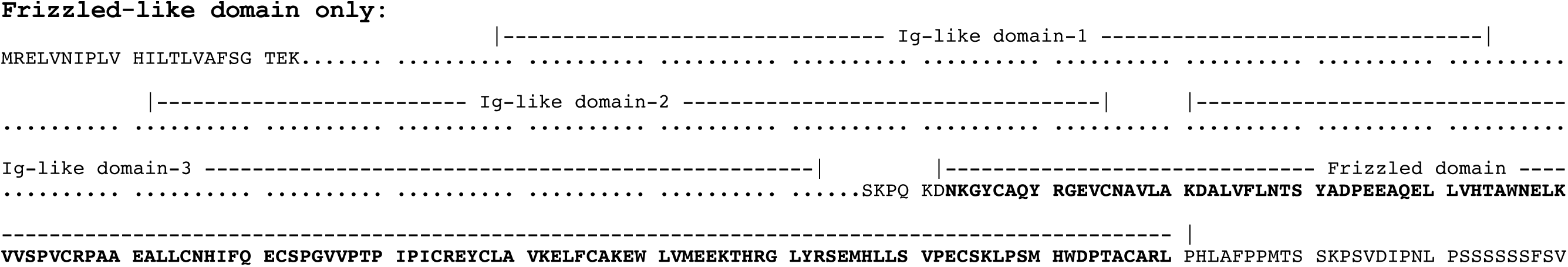
Protein sequences (amino acid) of different MuSK domain expression constructs. Bold letters indicate particular domains. Dots represent deleted regions.

**Figure S4.**
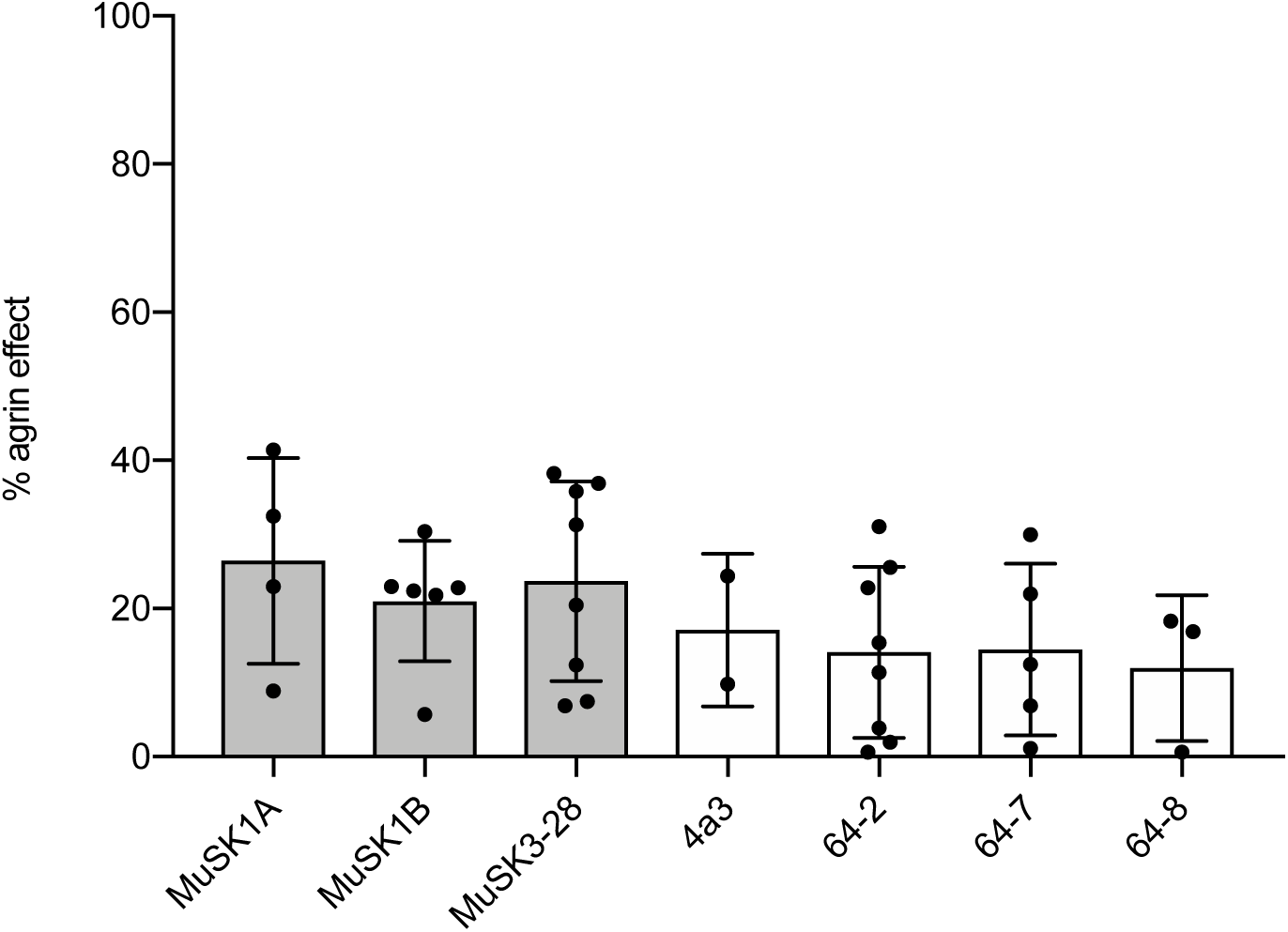
Antibody-induced AChR clustering in C2C12 mouse myotubes. The presence of agrin in C2C12 myotube cultures leads to dense clustering of AChRs that can be readily visualized with fluorescent α-bungarotoxin and quantified. Pathogenic MuSK autoantibodies can disrupt this clustering (*see main text and figures*). Three different human MuSK-specific mAbs (MuSK1A, MuSK1B, and MuSK3-28), the humanized murine control MuSK mAb 4A3 and three human non-MuSK-specific mAbs (64-2, 64-7, 64-8) were tested for their ability to induce the AChR clustering in the absence of agrin. Clustering of AChR was quantified relative to the measured effect of agrin. Quantitative results are normalized to agrin-only induced clustering. Each mAb was added to the cultures at 1μg/mL. Each data point represents the mean value from an independent experiment. Bars represent the mean of means and error bars SDs. Multiple comparisons ANOVA (against agrin), Dunnet’s test; * p<0.05, ** p<0.01, *** p<0.001, **** p<0.0001, only shown when significant).

## References

1. Vincent, A. 2002. Unravelling the pathogenesis of myasthenia gravis. Nat Rev Immunol 2: 797–804.

2. Gilhus, N. E. 2016. Myasthenia Gravis. N Engl J Med 375: 2570–2581.

3. Yi, J. S., J. T. Guptill, P. Stathopoulos, R. J. Nowak, and K. C. O’Connor. 2018. B cells in the pathophysiology of myasthenia gravis. Muscle Nerve 57: 172–184.

4. Vincent, A., D. Beeson, and B. Lang. 2000. Molecular targets for autoimmune and genetic disorders of neuromuscular transmission. Eur J Biochem 267: 6717–6728.

5. Lindstrom, J. M., A. G. Engel, M. E. Seybold, V. A. Lennon, and E. H. Lambert. 1976. Pathological mechanisms in experimental autoimmune myasthenia gravis. II. Passive transfer of experimental autoimmune myasthenia gravis in rats with anti-acetylcholine recepotr antibodies. J Exp Med 144: 739–753.

6. Shen, C., Y. Lu, B. Zhang, D. Figueiredo, J. Bean, J. Jung, H. Wu, A. Barik, D. M. Yin, W. C. Xiong, and L. Mei. 2013. Antibodies against low-density lipoprotein receptor-related protein 4 induce myasthenia gravis. J Clin Invest 123: 5190–5202.

7. Hoch, W., J. McConville, S. Helms, J. Newsom-Davis, A. Melms, and A. Vincent. 2001. Auto-antibodies to the receptor tyrosine kinase MuSK in patients with myasthenia gravis without acetylcholine receptor antibodies. Nat Med 7: 365–368.

8. Drachman, D. B., C. W. Angus, R. N. Adams, J. D. Michelson, and G. J. Hoffman. 1978. Myasthenic antibodies cross-link acetylcholine receptors to accelerate degradation. N Engl J Med 298: 1116–1122.

9. Zisimopoulou, P., P. Evangelakou, J. Tzartos, K. Lazaridis, V. Zouvelou, R. Mantegazza, C. Antozzi, F. Andreetta, A. Evoli, F. Deymeer, G. Saruhan-Direskeneli, H. Durmus, T. Brenner, A. Vaknin, S. Berrih-Aknin, M. Frenkian Cuvelier, T. Stojkovic, M. DeBaets, M. Losen, P. Martinez-Martinez, K. A. Kleopa, E. Zamba-Papanicolaou, T. Kyriakides, A. Kostera-Pruszczyk, P. Szczudlik, B. Szyluk, D. Lavrnic, I. Basta, S. Peric, C. Tallaksen, A. Maniaol, and S. J. Tzartos. 2014. A comprehensive analysis of the epidemiology and clinical characteristics of anti-LRP4 in myasthenia gravis. J Autoimmun 52: 139–145.

10. Higuchi, O., J. Hamuro, M. Motomura, and Y. Yamanashi. 2011. Autoantibodies to low-density lipoprotein receptor-related protein 4 in myasthenia gravis. Ann Neurol 69: 418–422.

11. Keung, B., K. R. Robeson, D. B. DiCapua, J. B. Rosen, K. C. O’Connor, J. M. Goldstein, and R. J. Nowak. 2013. Long-term benefit of rituximab in MuSK autoantibody myasthenia gravis patients. J Neurol Neurosurg Psychiatry 84: 1407–1409.

12. Huijbers, M. G., E. H. Niks, R. Klooster, M. de Visser, J. B. Kuks, J. H. Veldink, P. Klarenbeek, P. Van Damme, M. H. de Baets, S. M. van der Maarel, L. H. van den Berg, and J. J. Verschuuren. 2016. Myasthenia gravis with muscle specific kinase antibodies mimicking amyotrophic lateral sclerosis. Neuromuscul Disord 26: 350–353.

13. Nikolic, A. V., G. G. Bacic, M. Z. Dakovic, S. D. Lavrnic, V. M. Rakocevic Stojanovic, I. Z. Basta, and D. V. Lavrnic. 2015. Myopathy, muscle atrophy and tongue lipid composition in MuSK myasthenia gravis. Acta Neurol Belg 115: 361–365.

14. McConville, J., M. E. Farrugia, D. Beeson, U. Kishore, R. Metcalfe, J. Newsom-Davis, and A. Vincent. 2004. Detection and characterization of MuSK antibodies in seronegative myasthenia gravis. Ann Neurol 55: 580–584.

15. Aalberse, R. C., S. O. Stapel, J. Schuurman, and T. Rispens. 2009. Immunoglobulin G4: an odd antibody. Clin Exp Allergy 39: 469–477.

16. Klooster, R., J. J. Plomp, M. G. Huijbers, E. H. Niks, K. R. Straasheijm, F. J. Detmers, P. W. Hermans, K. Sleijpen, A. Verrips, M. Losen, P. Martinez-Martinez, M. H. De Baets, S. M. van der Maarel, and J. J. Verschuuren. 2012. Muscle-specific kinase myasthenia gravis IgG4 autoantibodies cause severe neuromuscular junction dysfunction in mice. Brain 135: 1081–1101.

17. Viegas, S., L. Jacobson, P. Waters, J. Cossins, S. Jacob, M. I. Leite, R. Webster, and A. Vincent. 2012. Passive and active immunization models of MuSK-Ab positive myasthenia: electrophysiological evidence for pre and postsynaptic defects. Exp Neurol 234: 506–512.

18. Shigemoto, K., S. Kubo, N. Maruyama, N. Hato, H. Yamada, C. Jie, N. Kobayashi, K. Mominoki, Y. Abe, N. Ueda, and S. Matsuda. 2006. Induction of myasthenia by immunization against muscle-specific kinase. J Clin Invest 116: 1016–1024.

19. Hopf, C., and W. Hoch. 1998. Dimerization of the muscle-specific kinase induces tyrosine phosphorylation of acetylcholine receptors and their aggregation on the surface of myotubes. J Biol Chem 273: 6467–6473.

20. Huijbers, M. G., W. Zhang, R. Klooster, E. H. Niks, M. B. Friese, K. R. Straasheijm, P. E. Thijssen, H. Vrolijk, J. J. Plomp, P. Vogels, M. Losen, S. M. Van der Maarel, S. J. Burden, and J. J. Verschuuren. 2013. MuSK IgG4 autoantibodies cause myasthenia gravis by inhibiting binding between MuSK and Lrp4. Proc Natl Acad Sci U S A 110: 20783–20788.

21. Koneczny, I., J. Cossins, P. Waters, D. Beeson, and A. Vincent. 2013. MuSK myasthenia gravis IgG4 disrupts the interaction of LRP4 with MuSK but both IgG4 and IgG1-3 can disperse preformed agrin-independent AChR clusters. PLoS One 8: e80695.

22. Otsuka, K., M. Ito, B. Ohkawara, A. Masuda, Y. Kawakami, K. Sahashi, H. Nishida, N. Mabuchi, A. Takano, A. G. Engel, and K. Ohno. 2015. Collagen Q and anti-MuSK autoantibody competitively suppress agrin/LRP4/MuSK signaling. Sci Rep 5: 13928.

23. Koneczny, I., J. A. Stevens, A. De Rosa, S. Huda, M. G. Huijbers, A. Saxena, M. Maestri, K. Lazaridis, P. Zisimopoulou, S. Tzartos, J. Verschuuren, S. M. van der Maarel, P. van Damme, M. H. De Baets, P. C. Molenaar, A. Vincent, R. Ricciardi, P. Martinez-Martinez, and M. Losen. 2017. IgG4 autoantibodies against muscle-specific kinase undergo Fab-arm exchange in myasthenia gravis patients. J Autoimmun 77: 104–115.

24. van der Neut Kolfschoten, M., J. Schuurman, M. Losen, W. K. Bleeker, P. Martinez-Martinez, E. Vermeulen, T. H. den Bleker, L. Wiegman, T. Vink, L. A. Aarden, M. H. De Baets, J. G. van de Winkel, R. C. Aalberse, and P. W. Parren. 2007. Anti-inflammatory activity of human IgG4 antibodies by dynamic Fab arm exchange. Science 317: 1554–1557.

25. Losen, M., A. F. Labrijn, V. H. van Kranen-Mastenbroek, M. L. Janmaat, K. G. Haanstra, F. J. Beurskens, T. Vink, M. Jonker, B. A. t Hart, M. Mane-Damas, P. C. Molenaar, P. Martinez-Martinez, E. van der Esch, J. Schuurman, M. H. de Baets, and P. Parren. 2017. Hinge-deleted IgG4 blocker therapy for acetylcholine receptor myasthenia gravis in rhesus monkeys. Sci Rep 7: 992.

26. Diaz-Manera, J., E. Martinez-Hernandez, L. Querol, R. Klooster, R. Rojas-Garcia, X. Suarez-Calvet, J. L. Munoz-Blanco, C. Mazia, K. R. Straasheijm, E. Gallardo, C. Juarez, J. J. Verschuuren, and I. Illa. 2012. Long-lasting treatment effect of rituximab in MuSK myasthenia. Neurology 78: 189–193.

27. Nowak, R. J., D. B. Dicapua, N. Zebardast, and J. M. Goldstein. 2011. Response of patients with refractory myasthenia gravis to rituximab: a retrospective study. Ther Adv Neurol Disord 4: 259–266.

28. Mei, H. E., I. Wirries, D. Frolich, M. Brisslert, C. Giesecke, J. R. Grun, T. Alexander, S. Schmidt, K. Luda, A. A. Kuhl, R. Engelmann, M. Durr, T. Scheel, M. Bokarewa, C. Perka, A. Radbruch, and T. Dorner. 2015. A unique population of IgG-expressing plasma cells lacking CD19 is enriched in human bone marrow. Blood 125: 1739–1748.

29. Cambridge, G., M. J. Leandro, J. C. Edwards, M. R. Ehrenstein, M. Salden, M. Bodman-Smith, and A. D. Webster. 2003. Serologic changes following B lymphocyte depletion therapy for rheumatoid arthritis. Arthritis Rheum 48: 2146–2154.

30. Hall, R. P., 3rd, R. D. Streilein, D. L. Hannah, P. D. McNair, J. A. Fairley, A. Ronaghy, K. D. Edhegard, and M. C. Levesque. 2013. Association of serum B-cell activating factor level and proportion of memory and transitional B cells with clinical response after rituximab treatment of bullous pemphigoid patients. J Invest Dermatol 133: 2786–2788.

31. Stathopoulos, P., A. Kumar, J. A. V. Heiden, E. Pascual-Goni, R. J. Nowak, and K. C. O’Connor. 2018. Mechanisms underlying B cell immune dysregulation and autoantibody production in MuSK myasthenia gravis. Ann N Y Acad Sci 1412: 154–165.

32. Stathopoulos, P., A. Kumar, R. J. Nowak, and K. C. O’Connor. 2017. Autoantibody-producing plasmablasts after B cell depletion identified in muscle-specific kinase myasthenia gravis. JCI Insight 2: e94263–e94275.

33. Franz, B., K. F. May, Jr., G. Dranoff, and K. Wucherpfennig. 2011. Ex vivo characterization and isolation of rare memory B cells with antigen tetramers. Blood 118: 348–357.

34. Lee, J. Y., P. Stathopoulos, S. Gupta, J. M. Bannock, R. J. Barohn, E. Cotzomi, M. M. Dimachkie, L. Jacobson, C. S. Lee, H. Morbach, L. Querol, J. L. Shan, J. A. Vander Heiden, P. Waters, A. Vincent, R. J. Nowak, and K. C. O’Connor. 2016. Compromised fidelity of B-cell tolerance checkpoints in AChR and MuSK myasthenia gravis. Ann Clin Transl Neurol 3: 443–454.

35. Linnington, C., M. Webb, and P. L. Woodhams. 1984. A novel myelin-associated glycoprotein defined by a mouse monoclonal antibody. J Neuroimmunol 6: 387–396.

36. Owens, G. P., J. L. Bennett, H. Lassmann, K. C. O’Connor, A. M. Ritchie, A. Shearer, C. Lam, X. Yu, M. Birlea, C. DuPree, R. A. Williamson, D. A. Hafler, M. P. Burgoon, and D. Gilden. 2009. Antibodies produced by clonally expanded plasma cells in multiple sclerosis cerebrospinal fluid. Ann Neurol 65: 639–649.

37. Graus, Y. F., M. H. de Baets, P. W. Parren, S. Berrih-Aknin, J. Wokke, P. J. van Breda Vriesman, and D. R. Burton. 1997. Human anti-nicotinic acetylcholine receptor recombinant Fab fragments isolated from thymus-derived phage display libraries from myasthenia gravis patients reflect predominant specificities in serum and block the action of pathogenic serum antibodies. J Immunol 158: 1919–1929.

38. Ray, A., A. A. Amato, E. M. Bradshaw, K. J. Felice, D. B. DiCapua, J. M. Goldstein, I. E. Lundberg, R. J. Nowak, H. L. Ploegh, E. Spooner, Q. Wu, S. N. Willis, and K. C. O’Connor. 2012. Autoantibodies produced at the site of tissue damage provide evidence of humoral autoimmunity in inclusion body myositis. PLoS One 7: e46709.

39. Maillette de Buy Wenniger, L. J., M. E. Doorenspleet, P. L. Klarenbeek, J. Verheij, F. Baas, R. P. Elferink, P. P. Tak, N. de Vries, and U. Beuers. 2013. Immunoglobulin G4+ clones identified by next-generation sequencing dominate the B cell receptor repertoire in immunoglobulin G4 associated cholangitis. Hepatology 57: 2390–2398.

40. Schanz, M., T. Liechti, O. Zagordi, E. Miho, S. T. Reddy, H. F. Gunthard, A. Trkola, and M. Huber. 2014. High-throughput sequencing of human immunoglobulin variable regions with subtype identification. PLoS One 9: e111726.

41. Leite, M. I., S. Jacob, S. Viegas, J. Cossins, L. Clover, B. P. Morgan, D. Beeson, N. Willcox, and A. Vincent. 2008. IgG1 antibodies to acetylcholine receptors in ’seronegative’ myasthenia gravis. Brain 131: 1940–1952.

42. Waters, P., M. Woodhall, K. C. O’Connor, M. Reindl, B. Lang, D. K. Sato, M. Jurynczyk, G. Tackley, J. Rocha, T. Takahashi, T. Misu, I. Nakashima, J. Palace, K. Fujihara, M. I. Leite, and A. Vincent. 2015. MOG cell-based assay detects non-MS patients with inflammatory neurologic disease. Neurol Neuroimmunol Neuroinflamm 2: e89.

43. Brochet, X., M. P. Lefranc, and V. Giudicelli. 2008. IMGT/V-QUEST: the highly customized and integrated system for IG and TR standardized V-J and V-D-J sequence analysis. Nucleic Acids Res 36: W503–508.

44. Lefranc, M. P. 2001. IMGT, the international ImMunoGeneTics database. Nucleic Acids Res 29: 207–209.

45. Tse, N., M. Morsch, N. Ghazanfari, L. Cole, A. Visvanathan, C. Leamey, and W. D. Phillips. 2014. The neuromuscular junction: measuring synapse size, fragmentation and changes in synaptic protein density using confocal fluorescence microscopy. J Vis Exp.

46. Gomez, A. M., J. A. Stevens, M. Mane-Damas, P. Molenaar, H. Duimel, F. Verheyen, J. Cossins, D. Beeson, M. H. De Baets, M. Losen, and P. Martinez-Martinez. 2016. Silencing of Dok-7 in Adult Rat Muscle Increases Susceptibility to Passive Transfer Myasthenia Gravis. Am J Pathol 186: 2559–2568.

47. Evoli, A., P. A. Tonali, L. Padua, M. L. Monaco, F. Scuderi, A. P. Batocchi, M. Marino, and E. Bartoccioni. 2003. Clinical correlates with anti-MuSK antibodies in generalized seronegative myasthenia gravis. Brain 126: 2304–2311.

48. Deymeer, F., O. Gungor-Tuncer, V. Yilmaz, Y. Parman, P. Serdaroglu, C. Ozdemir, A. Vincent, and G. Saruhan-Direskeneli. 2007. Clinical comparison of anti-MuSK-vs anti-AChR-positive and seronegative myasthenia gravis. Neurology 68: 609–611.

49. Sanders, D. B., K. El-Salem, J. M. Massey, J. McConville, and A. Vincent. 2003. Clinical aspects of MuSK antibody positive seronegative MG. Neurology 60: 1978–1980.

50. Clements, C. S., H. H. Reid, T. Beddoe, F. E. Tynan, M. A. Perugini, T. G. Johns, C. C. Bernard, and J. Rossjohn. 2003. The crystal structure of myelin oligodendrocyte glycoprotein, a key autoantigen in multiple sclerosis. Proc Natl Acad Sci U S A 100: 11059–11064.

51. Stiegler, A. L., S. J. Burden, and S. R. Hubbard. 2006. Crystal structure of the agrin-responsive immunoglobulin-like domains 1 and 2 of the receptor tyrosine kinase MuSK. J Mol Biol 364: 424–433.

52. Stiegler, A. L., S. J. Burden, and S. R. Hubbard. 2009. Crystal structure of the frizzled-like cysteine-rich domain of the receptor tyrosine kinase MuSK. J Mol Biol 393: 1–9.

53. Huijbers, M. G., A. F. Vink, E. H. Niks, R. H. Westhuis, E. W. van Zwet, R. H. de Meel, R. Rojas-Garcia, J. Diaz-Manera, J. B. Kuks, R. Klooster, K. Straasheijm, A. Evoli, I. Illa, S. M. van der Maarel, and J. J. Verschuuren. 2016. Longitudinal epitope mapping in MuSK myasthenia gravis: implications for disease severity. J Neuroimmunol 291: 82–88.

54. Kim, N., A. L. Stiegler, T. O. Cameron, P. T. Hallock, A. M. Gomez, J. H. Huang, S. R. Hubbard, M. L. Dustin, and S. J. Burden. 2008. Lrp4 is a receptor for Agrin and forms a complex with MuSK. Cell 135: 334–342.

55. Cardona, A., O. Pritsch, G. Dumas, J. F. Bach, and G. Dighiero. 1995. Evidence for an antigen-driven selection process in human autoantibodies against acetylcholine receptor. Mol Immunol 32: 1215–1223.

56. Farrar, J., S. Portolano, N. Willcox, A. Vincent, L. Jacobson, J. Newsom-Davis, B. Rapoport, and S. M. McLachlan. 1997. Diverse Fab specific for acetylcholine receptor epitopes from a myasthenia gravis thymus combinatorial library. Int Immunol 9: 1311–1318.

57. Sims, G. P., H. Shiono, N. Willcox, and D. I. Stott. 2001. Somatic hypermutation and selection of B cells in thymic germinal centers responding to acetylcholine receptor in myasthenia gravis. J Immunol 167: 1935–1944.

58. Saxena, A., J. Stevens, H. Cetin, I. Koneczny, R. Webster, K. Lazaridis, S. Tzartos, K. Vrolix, G. Nogales-Gadea, B. Machiels, P. C. Molenaar, J. Damoiseaux, M. H. De Baets, K. Simon-Keller, A. Marx, A. Vincent, M. Losen, and P. Martinez-Martinez. 2017. Characterization of an anti-fetal AChR monoclonal antibody isolated from a myasthenia gravis patient. Sci Rep 7: 14426.

59. Vrolix, K., J. Fraussen, M. Losen, J. Stevens, K. Lazaridis, P. C. Molenaar, V. Somers, M. A. Bracho, R. Le Panse, P. Stinissen, S. Berrih-Aknin, J. G. Maessen, L. Van Garsse, W. A. Buurman, S. J. Tzartos, M. H. De Baets, and P. Martinez-Martinez. 2014. Clonal heterogeneity of thymic B cells from early-onset myasthenia gravis patients with antibodies against the acetylcholine receptor. J Autoimmun 52: 101–112.

60. Bennett, J. L., C. Lam, S. R. Kalluri, P. Saikali, K. Bautista, C. Dupree, M. Glogowska, D. Case, J. P. Antel, G. P. Owens, D. Gilden, S. Nessler, C. Stadelmann, and B. Hemmer. 2009. Intrathecal pathogenic anti-aquaporin-4 antibodies in early neuromyelitis optica. Ann Neurol 66: 617–629.

61. Colliou, N., D. Picard, F. Caillot, S. Calbo, S. Le Corre, A. Lim, B. Lemercier, B. Le Mauff, M. Maho-Vaillant, S. Jacquot, C. Bedane, P. Bernard, F. Caux, C. Prost, E. Delaporte, M. S. Doutre, B. Dreno, N. Franck, S. Ingen-Housz-Oro, O. Chosidow, C. Pauwels, C. Picard, J. C. Roujeau, M. Sigal, E. Tancrede-Bohin, I. Templier, R. Eming, M. Hertl, M. D’Incan, P. Joly, and P. Musette. 2013. Long-term remissions of severe pemphigus after rituximab therapy are associated with prolonged failure of desmoglein B cell response. Sci Transl Med 5: 175ra130.

62. Wilson, R., M. Makuch, A. K. Kienzler, J. Varley, J. Taylor, M. Woodhall, J. Palace, M. I. Leite, P. Waters, and S. R. Irani. 2018. Condition-dependent generation of aquaporin-4 antibodies from circulating B cells in neuromyelitis optica. Brain 141: 1063–1074.

63. Makuch, M., R. Wilson, A. Al-Diwani, J. Varley, A. K. Kienzler, J. Taylor, A. Berretta, D. Fowler, B. Lennox, M. I. Leite, P. Waters, and S. R. Irani. 2018. N-methyl-D-aspartate receptor antibody production from germinal center reactions: Therapeutic implications. Ann Neurol 83: 553–561.

64. Di Zenzo, G., G. Di Lullo, D. Corti, V. Calabresi, A. Sinistro, F. Vanzetta, B. Didona, G. Cianchini, M. Hertl, R. Eming, M. Amagai, B. Ohyama, T. Hashimoto, J. Sloostra, F. Sallusto, G. Zambruno, and A. Lanzavecchia. 2012. Pemphigus autoantibodies generated through somatic mutations target the desmoglein-3 cis-interface. J Clin Invest 122: 3781–3790.

65. Cho, M. J., A. S. Lo, X. Mao, A. R. Nagler, C. T. Ellebrecht, E. M. Mukherjee, C. M. Hammers, E. J. Choi, P. M. Sharma, M. Uduman, H. Li, A. H. Rux, S. A. Farber, C. B. Rubin, S. H. Kleinstein, B. S. Sachais, M. R. Posner, L. A. Cavacini, and A. S. Payne. 2014. Shared VH1-46 gene usage by pemphigus vulgaris autoantibodies indicates common humoral immune responses among patients. Nat Commun 5: 4167.

66. Makino, T., R. Nakamura, M. Terakawa, S. Muneoka, K. Nagahira, Y. Nagane, J. Yamate, M. Motomura, and K. Utsugisawa. 2017. Analysis of peripheral B cells and autoantibodies against the anti-nicotinic acetylcholine receptor derived from patients with myasthenia gravis using single-cell manipulation tools. PLoS One 12: e0185976.

67. Rodriguez Cruz, P. M., S. Huda, P. Lopez-Ruiz, and A. Vincent. 2015. Use of cell-based assays in myasthenia gravis and other antibody-mediated diseases. Exp Neurol 270: 66–71.

68. Basta, I., A. Nikolic, M. Losen, P. Martinez-Martinez, V. Stojanovic, S. Lavrnic, M. de Baets, and D. Lavrnic. 2012. MuSK myasthenia gravis and Lambert-Eaton myasthenic syndrome in the same patient. Clin Neurol Neurosurg 114: 795–797.

69. Oh, S. J. 2016. Myasthenia gravis Lambert-Eaton overlap syndrome. Muscle Nerve 53: 20–26.

70. Tsonis, A. I., P. Zisimopoulou, K. Lazaridis, J. Tzartos, E. Matsigkou, V. Zouvelou, R. Mantegazza, C. Antozzi, F. Andreetta, A. Evoli, F. Deymeer, G. Saruhan-Direskeneli, H. Durmus, T. Brenner, A. Vaknin, S. Berrih-Aknin, A. Behin, T. Sharshar, M. De Baets, M. Losen, P. Martinez-Martinez, K. A. Kleopa, E. Zamba-Papanicolaou, T. Kyriakides, A. Kostera-Pruszczyk, P. Szczudlik, B. Szyluk, D. Lavrnic, I. Basta, S. Peric, C. Tallaksen, A. Maniaol, C. Casasnovas Pons, J. Pitha, M. Jakubikova, F. Hanisch, and S. J. Tzartos. 2015. MuSK autoantibodies in myasthenia gravis detected by cell based assay--A multinational study. J Neuroimmunol 284: 10–17.

71. Cho, A., B. Bradley, R. Kauffman, L. Priyamvada, Y. Kovalenkov, R. Feldman, and J. Wrammert. 2017. Robust memory responses against influenza vaccination in pemphigus patients previously treated with rituximab. JCI Insight 2.

72. Niks, E. H., Y. van Leeuwen, M. I. Leite, F. W. Dekker, A. R. Wintzen, P. W. Wirtz, A. Vincent, M. J. van Tol, C. M. Jol-van der Zijde, and J. J. Verschuuren. 2008. Clinical fluctuations in MuSK myasthenia gravis are related to antigen-specific IgG4 instead of IgG1. J Neuroimmunol 195: 151–156.

73. Zhang, W., A. S. Coldefy, S. R. Hubbard, and S. J. Burden. 2011. Agrin binds to the N-terminal region of Lrp4 protein and stimulates association between Lrp4 and the first immunoglobulin-like domain in muscle-specific kinase (MuSK). J Biol Chem 286: 40624–40630.

74. Mori, S., S. Yamada, S. Kubo, J. Chen, S. Matsuda, M. Shudou, N. Maruyama, and K. Shigemoto. 2012. Divalent and monovalent autoantibodies cause dysfunction of MuSK by distinct mechanisms in a rabbit model of myasthenia gravis. J Neuroimmunol 244: 1–7.

75. Cantor, S., W. Zhang, N. Delestree, L. Remedio, G. Z. Mentis, and S. J. Burden. 2018. Preserving neuromuscular synapses in ALS by stimulating MuSK with a therapeutic agonist antibody. Elife 7.

76. Kowarik, M. C., D. Astling, C. Gasperi, S. Wemlinger, H. Schumann, M. Dzieciatkowska, A. M. Ritchie, B. Hemmer, G. P. Owens, and J. L. Bennett. 2017. CNS Aquaporin-4-specific B cells connect with multiple B-cell compartments in neuromyelitis optica spectrum disorder. Ann Clin Transl Neurol 4: 369–380.

77. Chen, J., Q. Zheng, C. M. Hammers, C. T. Ellebrecht, E. M. Mukherjee, H. Y. Tang, C. Lin, H. Yuan, M. Pan, J. Langenhan, L. Komorowski, D. L. Siegel, A. S. Payne, and J. R. Stanley. 2017. Proteomic Analysis of Pemphigus Autoantibodies Indicates a Larger, More Diverse, and More Dynamic Repertoire than Determined by B Cell Genetics. Cell Rep 18: 237–247.

78. Kasperkiewicz, M., C. T. Ellebrecht, H. Takahashi, J. Yamagami, D. Zillikens, A. S. Payne, and M. Amagai. 2017. Pemphigus. Nat Rev Dis Primers 3: 17026.

79. Querol, L., J. Devaux, R. Rojas-Garcia, and I. Illa. 2017. Autoantibodies in chronic inflammatory neuropathies: diagnostic and therapeutic implications. Nat Rev Neurol 13: 533–547.

